# Male sex chromosomal complement exacerbates the pathogenicity of Th17 cells in a chronic model of CNS autoimmunity

**DOI:** 10.1101/640904

**Authors:** Prenitha Mercy Ignatius Arokia Doss, Joanie Baillargeon, Asmita Pradeep Yeola, John B. Williams, Mickael Leclercq, Charles Joly-Beauparlant, Melanie Alpaugh, Ana C. Anderson, Arnaud Droit, Hans Lassmann, Craig S. Moore, Manu Rangachari

## Abstract

Sex differences in the incidence and severity of multiple sclerosis (MS) have long been recognized. However, the underlying cellular and molecular mechanisms for why male sex is associated with more aggressive and debilitating disease remain poorly defined. Using an T cell adoptive transfer model of chronic EAE, we find that male Th17 cells induced disease of increased severity relative to female Th17 cells, irrespective of whether transferred to male or female recipients. Throughout the disease course, a greater frequency of male Th17 cells produced the heterodox cytokine IFNγ, a hallmark of pathogenic Th17 responses. Intriguingly, sex chromosomal complement, and not hormones, were responsible for the increased pathogenicity of male Th17 cells and an X-linked immune regulator, *Jarid1c*, was downregulated in both pathogenic male Th17 and CD4^+^ T cells from men with MS. Together, our data indicate that male sex critical regulates Th17 cell plasticity and pathogenicity via sex chromosomal complement.

## Introduction

Multiple sclerosis (MS) is a chronic inflammatory disease in which T cells breach the blood-brain barrier and mount an attack against central nervous system (CNS) myelin. MS, which affects more than 2 million people world-wide (Browne et al., 2014), presents two broad clinical courses: relapsing/remitting (RR) or progressive. A number of immunomodulatory therapies are now available for RR disease, which is seen in approximately 80% of patients from onset (Scalfari et al., 2013). However, 30-60% of MS patients will ultimately transition to a secondary progressive (SP) form of the disease for which current drugs are largely ineffective (Larochelle et al., 2016).

The role of biological sex as a determinant of MS incidence is well established, as women develop the disease at 2-3 times the frequency of men (Greer and McCombe, 2011; Sloka et al., 2011). However, what is less well understood is why MS in men tends to progress more rapidly and aggressively (Beatty and Aupperle, 2002; Menon et al., 2013; Tremlett et al., 2006). In parallel, it is increasingly more appreciated that T helper 17 (Th17) CD4^+^ T cell responses may be drivers of progressive MS (Huber et al., 2014; Romme Christensen et al., 2013), though the mechanisms remain unclear. Together, these clinical findings illustrate the uncoupling of disease incidence and severity in MS, and underscore the need for deciphering the cellular and molecular mechanisms underlying the role of biological sex in determining disease course in MS.

Experimental autoimmune encephalomyelitis (EAE) is a mouse model that recapitulates key immune aspects of MS. It can be induced by active immunization by encephalitogenic myelin-derived peptides, as well by the adoptive transfer of primed myelin-reactive T cells, with the latter approach permitting one to isolate the role of T cells, and specific effector T cell subsets, in disease processes (Jäger et al., 2009; Rangachari et al., 2012). The popular C57BL6/J model of EAE displays either an acute monophasic or stably chronic disease course (Berard et al., 2010), while EAE in SJL/J mice shows a RR pattern (McRae et al., 1995). By contrast, numerous groups have shown that active immunization of non-obese diabetic (NOD) strain mice with myelin oligodendrocyte glycoprotein (MOG)_[35−55]_ induces a RR disease course followed by transition to chronic disease. Accordingly, the NOD-EAE active immunization model has been used to probe potential mechanisms driving progressive disease (Farez et al., 2009; Mayo et al., 2014).

In this study, we used a T cell receptor (TcR) transgenic (Tg) model (1C6), that recognizes MOG_[35−55]_ on the NOD background (Anderson et al., 2012), to examine the role of biological sex in determining Th17 phenotype and disease course upon adoptive transfer into NOD.*Scid* mice. We observed a striking sex bias, as recipients of male 1C6 Th17 cells developed disease of greater severity than recipients of female Th17 cells, characterized by a higher frequency of progressive disease. This sex dimorphism was T cell-intrinsic, as male Th17 cells induced severe disease irrespective of whether they were transferred to male or female recipients. Male Th17 cells displayed a strong tendency towards production of interferon (IFN)-γ upon adoptive transfer, and they developed an “ex-Th17” transcriptional signature that is associated with Th17 cell pathogenicity. Notably, male Th17 pathogenicity was driven my male sex chromosomal complement rather than by androgens; we identified an X-linked immunomodulatory gene, *Jarid1c*, which escapes second-copy silencing in female immune cells and is downregulated in both male 1C6 Th17 cells that transfer severe EAE. Interestingly, the downregulation of Jarid1c was also observed in CD4^+^ T cells taken from male patients with MS. We thus uncover a molecular and cellular basis for the association of male sex with exacerbated disease course in CNS autoimmunity.

## Results

### Male 1C6 Th17 cells induce severe and frequently lethal progressive EAE

Multiple lines of evidence indicate a role for dysregulated Th17 responses in progressive MS. CD4^+^IL-23R^+^ Th17 cells are increased in the peripheral blood of SPMS and primary progressive (PP) MS, but not RRMS, patients (Romme Christensen et al., 2013). Further, circulating levels of Th17-inducible chemokines are higher in SPMS as compared to RRMS (Huber et al., 2014). However, while MS progresses more rapidly in men, the potential contribution of biological sex to Th17-driven relapsing/remitting and chronic progressive CNS autoimmune inflammation has not been extensively studied.

To examine this, we developed an adoptive transfer protocol in which we first purified splenic and lymph node CD4^+^CD62L^hi^ cells from female or male 1C6 mice and stimulated them for 5 days with agonistic antibodies to CD3 and CD28 under established pathogenic Th17 differentiation conditions (Bettelli et al., 2006) (Figure 1A). By using an antibody stimulation approach, rather than stimulation via peptide and antigen-presenting cells, we were able to isolate the contributions of T cells alone to disease processes. After 5 days of *in vitro* culture, both female and male Th17 cells were highly positive for the Th17 master transcription factor RORγt (Figure 1B), indicating that they were committed to the Th17 lineage prior to *in vivo* transfer. We then adoptively transferred 1C6 Th17 cells to sex-matched NOD.*Scid* recipient mice and monitored them for signs of clinical disease over the course of 70 days. Use of NOD.*Scid* mice as hosts carried several distinct advantages. First, NOD.*Scid* are resistant to the development of spontaneous diabetes that is characteristic of the NOD strain (Bobbala et al., 2012; Christianson et al., 1993; Rohane et al., 1995) and thus we could exclude insulitis and beta-cell destruction as confounding factors that could affect the course of EAE. Further, the lack of endogenous T cells in NOD.*Scid* mice allowed us to conduct sex-mismatched transfers without fear of host-versus-graft reactivity.

**Figure 1.**
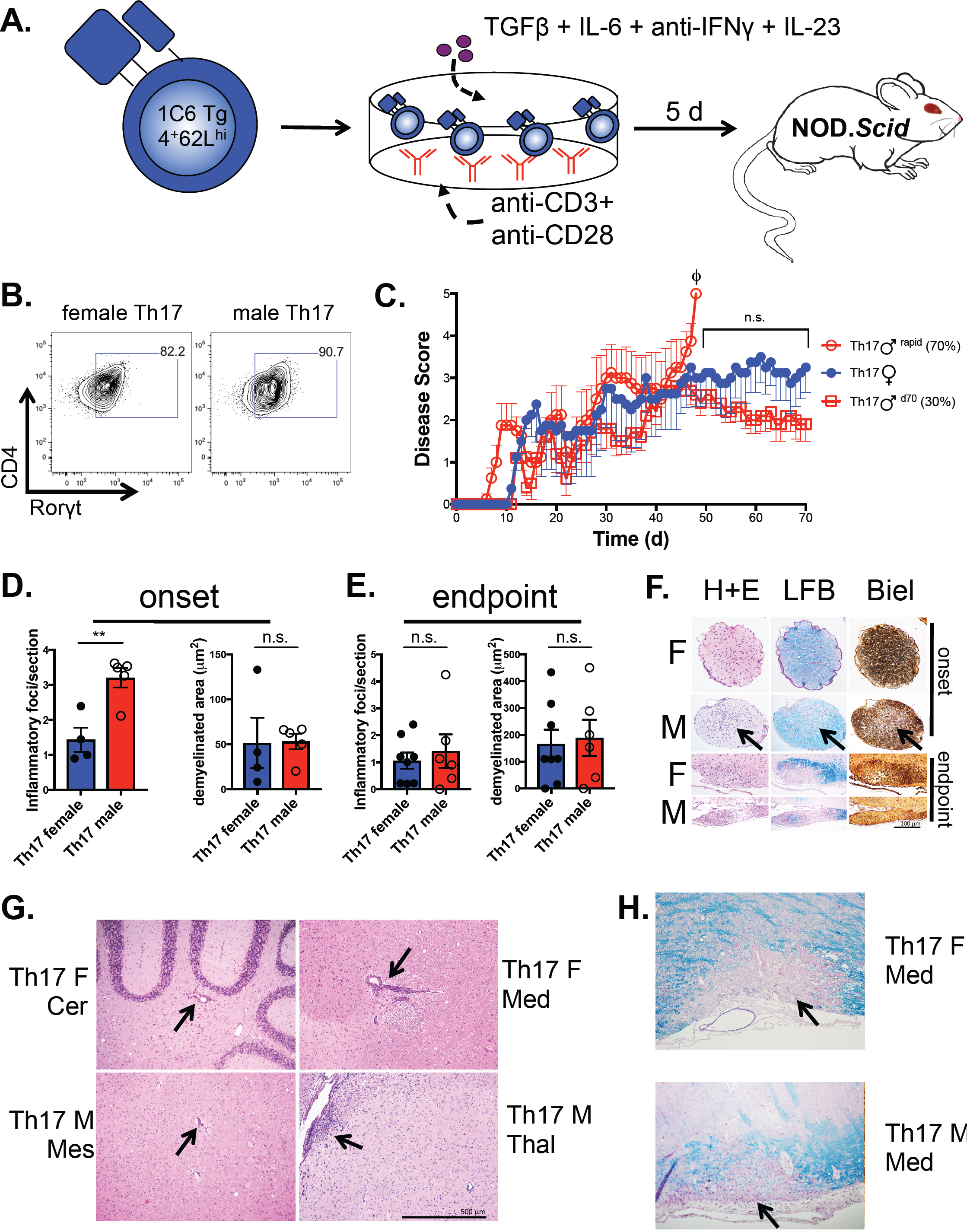
Male 1C6 Th17 transfer disease of earlier onset and exacerbated severity. **A.** Schematic depicting the experimental approach. Tg, transgenic. **B**. Female and male 1C6 Th17 cells were generated *in vitro* as depicted in (**A**) and Rorγt expression was measured by flow cytometry at d5 of culture prior to adoptive transfer. Gated on live CD4^+^ events. Representative of 3 experiments. **C.** Female and male 1C6 Th17 were adoptively transferred to sex-matched NOD.*Scid* recipients that were monitored for signs of EAE up to d70 post-transfer. A proportion of male recipients (M^rapid^, n=4) attained ethical endpoints before d70. No significant differences in day-to-day disease score were observed between males that survived to d70 (M^late^, n=5) and females. Φ, all mice in the group attained ethical endpoints. Representative of 4 transfer experiments. Percentages in the graph legend refer to all mice studied. **D-H**. Female or male 1C6 Th17 were transferred sex-matched NOD.*Scid* that were sacrificed at disease onset (**D, F, G**) or at experimental endpoints (**E, F, H**). Female, n=8; Male, n=6. All male endpoint mice experienced a rapid disease course; all female endpoint mice survived to d70. **D, E**. The presence of inflammatory foci (H&E) and demyelination (Luxol fast blue, LFB) were quantified (**D**) from paraffinized spinal cord sections at onset (4 females, 5 males) or endpoint (8 females, 6 males). Symbols represent individual mice. Error bars represent s.e.m. ** p<0.01, *t*-test. **F.** Optic nerve sections were taken from female and male recipients at onset or endpoint. Leukocyte infiltration (H&E), demyelination (LFB) and axon damage (Bielschowsky’s silver stain; Biel) were assessed. Representative of 4 females and 4 males. **G.** Inflammatory lesions in the CNS of Th17 recipients at disease onset. Clockwise from upper left: cerebellum (Cer), Th17 female; medulla (Med), Th17 female; thalamus (Thal), Th17 male; mesencephalon (Mes), Th17 male. **H.** Demyelinating lesions in the medullae of a female (*top*) and male (*bottom*) Th17 recipient at endpoint. 10X magnification for all images.

Strikingly, adoptive transfer of male 1C6 Th17 cells to male NOD.*Scid* induced EAE of much greater severity overall than that seen when female 1C6 Th17 were transferred to female NOD.*Scid* (Table 1). More than two-thirds of male Th17 recipients (M^rapid^) developed a chronically worsening disease course and rapidly attained ethical endpoints (33.7 d ± 2.5). The remaining males developed a non-lethal course (M^d70^) that did not differ in severity from female (F) Th17 recipients (Figure 1C, Table 1). All female Th17 recipients survived to d70. These data gave the first indication that male Th17 cells have a greater intrinsic pathogenic capacity.

**Table 1.** Clinical features of recipients of female or male 1C6 effector CD4^+^ T cells. All mice presented here were followed to experimental endpoints. The following groups were compared: 1:2, 1:2a:2b, 3:4, 2:4. ^a^ p<0.0001; ^b^ p<0.0001, ^c^ p=0.0094; ^d^ p=0.021; ^e^ p=0.0194; ^f^ p=0.0331; ^g^ p<0.0001; ^h^ p<0.0001; ^i^ p<0.0001;^j^ p<0.0001; ^k^ p=0.0012; ^l^ p=0.001; ^m^ p<0.0001; ^n^ p=0.0013; ^o^ p=0.0063; ^p^ p=0.0128. All other comparisons were not significant. Onset and number of relapses were assessed by *t*-test for pairwise comparisons or by Tukey’s multiple comparisons test after one-way ANOVA (1:2a:2b). Mean maximal score was assessed by Mann-Whitney *U* test, or by Dunn’s multiple comparisons test after one-way ANOVA (1:2a:2b). Incidence of overall disease, chronic progression, relapse-only, and relapse/chronic were assessed with Fisher’s exact test, with Bonferroni’s correction applied to 1:2a:2b.

| *Group #* | *Group* | *Incidence* | *Onset (days)* | *Mean maximal score* | *Number of relapses* | *Rapid course* | *Chronic progressive* | *Relapse only* | *Relapse/Chronic* |
| --- | --- | --- | --- | --- | --- | --- | --- | --- | --- |
| 1 | F (all)<br>Th17 | 30/30 | 12.4 ± 0.6 | 2.6 ± 0.2<sup>a,b</sup> | 1.2 ± 0.2<sup>e,f</sup> | 0/30<sup>h,i</sup> | 10/30<sup>l,m</sup> | 13/30 | 10/30 |
| 2 | M (all)<br>Th17 | 33/33 | 11.7 ± 0.6 | 4.3 ± 0.5<sup>a,d</sup> | 0.96 ± 0.2 | 23/33<sup>h,k</sup> | 25/33<sup>l</sup> | 11/33 | 9/33 |
| 2a | *M<sup>rapid</sup>*<br>*Th17* | 23/23 | 12.3 ± 0.8 | 5 ± 0<sup>b,c</sup> | 0.52 ± 0.1<sup>e,g</sup> | 23/23<sup>i,j</sup> | 20/23<sup>m</sup> | 6/23 | 4/23 |
| 2b | *M<sup>d70</sup>*<br>*Th17* | 10/10 | 10.7 ± 0.4 | 2.9 ± 0.1<sup>c</sup> | 2 ± 0.2<sup>f,g</sup> | 0/10<sup>j</sup> | 5/10 | 5/10 | 5/10 |
| 3 | F Th1 | 14/14 | 15.6± 2.0 | 2.4 ± 0.9 | 1.9 ± 0.3 | 0/14 | 1/14<sup>n</sup> | 12/14<sup>o</sup> | 1/14<sup>p</sup> |
| 4 | M Th1 | 13/13 | 12.8± 1.3 | 3.3 ± 0.2<sup>d</sup> | 1.7 ± 0.3 | 2/13<sup>k</sup> | 9/13<sup>n</sup> | 4/13<sup>o</sup> | 7/13<sup>p</sup> |

Irrespective of sex, more than half of all Th17 recipients (35/63) developed chronically worsening disease, while close to a third (19/63) displayed at least one relapse/remission cycle followed by chronic worsening of symptoms. Thus, a substantial frequency of 1C6 Th17 recipients recapitulated the biphasic disease course that is observed in actively immunized NOD animals. Further, as NOD.*Scid* mice do not develop diabetes, this also indicated that biphasic disease could develop independently of the insulitis and beta cell destruction that characterizes the NOD strain.

We next examined classical parameters of CNS damage to see whether they might help explain the increased severity of disease seen in the majority of male Th17 recipients. We observed a significant increase in leukocytic infiltration of the spinal cord at disease onset in males relative to females (Figure 1D), but no differences in area of demyelination. We observed no differences between male and female recipients at endpoint with respect to either inflammatory infiltration or demyelination in the spinal cord (Figure 1E), despite the fact that all the males analyzed had a rapid disease course; this was in line with previous findings that histopathological markers of CNS damage may be insensitive correlates of EAE severity (Goncalves DaSilva et al., 2010).

When NOD.*Scid* mice were reconstituted with naive 1C6 CD4^+^ T cells and then actively immunized, a significant frequency developed optic neuritis (Anderson et al., 2012). We therefore examined the effect of sex on the induction of optic neuritis. Strikingly, male Th17 mice displayed leukocytic inflammation, demyelination and axon degeneration in the optic nerve at disease onset, while female recipients showed no signs of histological damage (Figure 1F). In endstage disease, optic nerve damage was observed in all mice regardless of sex. Thus, male sex accelerates optic nerve damage, similar to its effects on paralytic disease course.

Actively immunized NOD mice display inflammatory lesions in both the cerebellum and midbrain (Dang et al., 2015). We next examined whether adoptive transfer of 1C6 Th17 could cause lesion formation in the brain as well as the spinal cord. Meningeal inflammation was observed in the brains of 100% of female (4/4) and male (5/5) recipients at onset, and in 75% of female (6/8) and 50% of male (3/6) mice at endpoint. We saw widespread evidence of brain parenchymal inflammation at onset (female, 100%, 4/4; male, 100% 5/5) and endpoint (female, 75%, 6/8; male, 83%, 5/6). Inflammatory lesions (Figure 1G) were frequently observed in the cerebellum, medulla, mesencephalon and cerebellum, and less frequently in the thalamus. Demyelinated lesions were never seen at onset, but were observed in the medullae of 2/8 females and 3/6 males at endpoint (Fig 1H). In sum, our data thus far showed that adoptive transfer of male 1C6 Th17 to male recipients could induce chronic EAE of greater severity than that observed in female recipients of female Th17. Further, disease in males was characterized by increased spinal cord inflammation and optic nerve damage are exacerbated at onset.

### Transferred 1C6 Th17 cells display phenotypic plasticity at early disease timepoints

We next addressed whether the severe disease seen in male Th17 recipients resulted from augmented pathogenicity of male Th17 cells themselves, or if it was caused by T cell-extrinsic factors such as increased sensitivity of the male CNS to inflammatory insult. In fact, recent studies indicated that increased susceptibility of male SJL mice to EAE is the result of increased neurodegeneration in the male CNS (Du et al., 2014). To address this question, we generated and adoptively transferred either male or female 1C6 Th17 cells to sex-matched or –mismatched recipients. Here, we exploited the fact that NOD.*Scid* do not have endogenous T cells and therefore would not reject sex-mismatched donor cells. Male 1C6 Th17 cells transferred far more severe disease relative to female Th17 cells, irrespective of whether transferred to female (Figure 2A) or male (Figure 2B) recipient NOD.*Scid* mice. Taken together, our data showed that male sex is a key cell intrinsic determinant of the severity of disease induced by Th17 cells.

**Figure 2.**
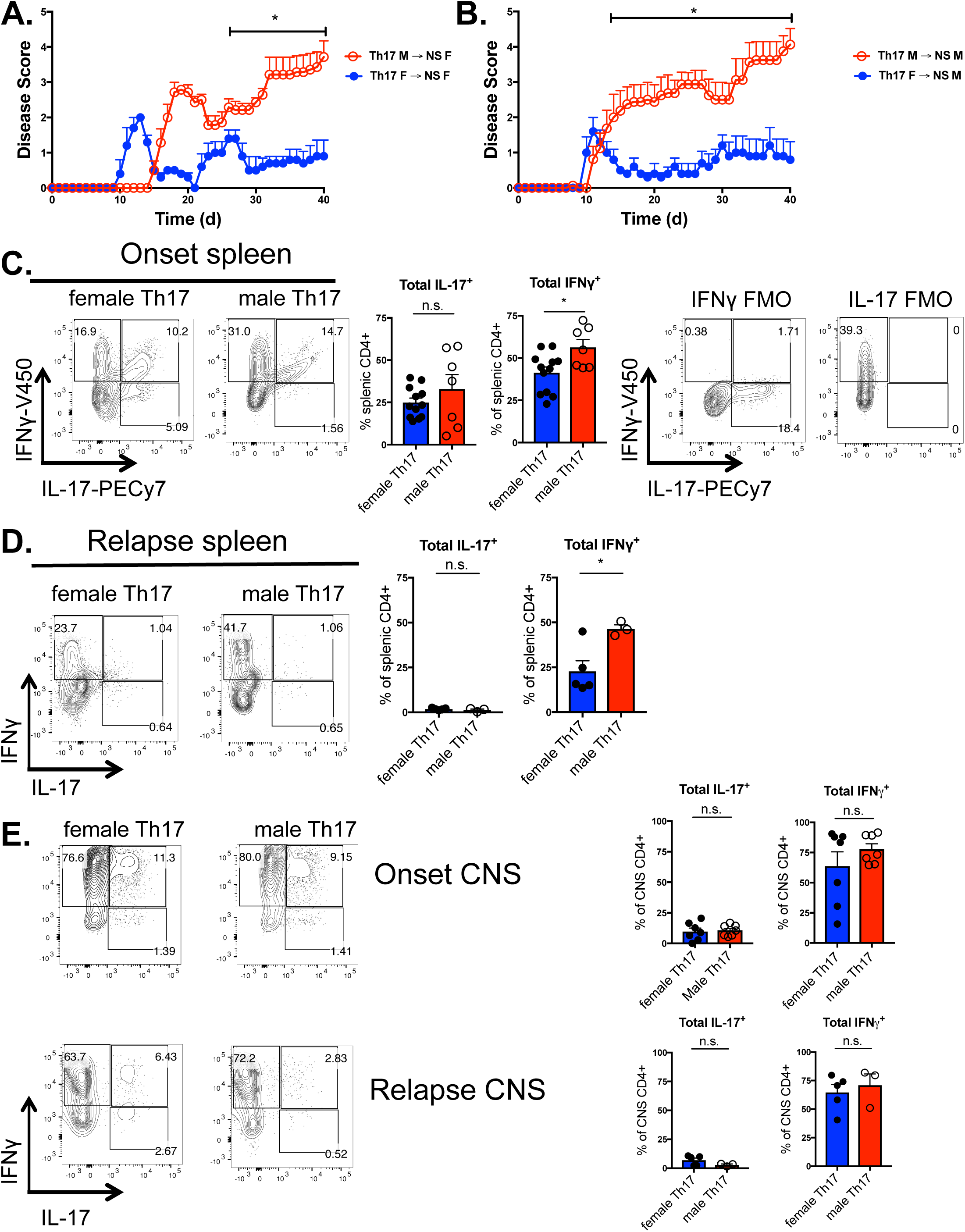
Peripheral male Th17 show increased IFNγ in early disease. **A.** Male or female 1C6 Th17 were transferred to female NOD.*Scid* that were monitored for signs of EAE. M→F, n=7; F→F, n=6. **B.** Male or female 1C6 Th17 were transferred to male NOD.*Scid* that were monitored for signs of EAE. M→M, n=8; F→M, n=5. * p<0.05, Mann-Whitney *U* test. **C**. Female recipients of female Th17 or male recipients of male Th17 were sacrificed at disease onset, and production of IL-17 and IFNγ from splenic CD4^+^ T cells was assessed by intracellular flow cytometry. FACS plots representative of F, n=12; M, n=7. * p<0.05, t-test. Rightmost FACS plots, fluorescence minus one (FMO) controls from the male mouse presented in this panel. **D.** Production of IL-17 and IFNγ from CD4^+^ splenic T cells from female or male Th17 cells at relapse. F, n=5; M, n=3. * p<0.05, *t*-test. **E.** Production of IL-17 and IFNγ from CNS-infiltrating CD4^+^ T cells from male or female Th17 recipients at onset (*top row*) or relapse (*bottom row*). Error bars, s.e.m. Each circle represents an individual mouse.

We next sought to understand why male Th17 cells could induce EAE of increased severity. Production of the signature cytokine IL-17A does not, on its own, necessarily dictate whether a Th17 cell is pathogenic (Y. Lee et al., 2012), and the IL-17^+^ Th17 phenotype is highly unstable (Hirota et al., 2011; Kurschus et al., 2010). Further, Th17 cells readily express the Th1 signature cytokine IFNγ *in vivo* (Gaublomme et al., 2015; Zhu et al., 2010), and these IFNγ^+^ Th17 cells have been implicated in the pathogenesis of EAE (Abromson-Leeman et al., 2009; Carbajal et al., 2015; Suryani and Sutton, 2007). It was therefore possible that an increase in phenotypic plasticity could explain the augmented pathogenicity of male Th17 cells.

To address this, we first isolated male versus female 1C6 Th17 cells from sex-matched NOD.*Scid* recipients at disease onset and measured their co-production of IFNγ and IL-17. Surprisingly, we detected few *bonafide* IL-17^+^IFNγ^−^ CD4^+^T cells of either sex; the vast majority of T cells either co-produced IL-17 and IFNγ, or they produced IFNγ alone (Fig 2C). Notably, while production of IL-17 did not differ between male and female Th17 cells, a significantly greater proportion of male Th17 cells produced IFNγ (Fig 2C). At the first relapse, IL-17 production was almost completely absent from both female and male cells in the spleen, indicating that production of this cytokine from T cells was not essential to pathology after the earliest stage of visible disease. Nevertheless, the percentage of IFNγ-producing male splenic cells was again significantly higher than that observed of females at relapse (Fig 2D). Thus, despite being committed to the Th17 lineage prior to adoptive transfer (Fig 1B), transferred Th17 cells, and male Th17 cells in particular rapidly exhibited phenotypic plasticity in the *in vivo* setting.

To examine whether sex-specific differences could be observed in the target organ, we next measured cytokine production of T cells entering the CNS at onset and relapse. At both timepoints, the vast majority of T cells, both male and female, produced no IL-17 and rather displayed an IL-17^−^IFNγ^+^ phenotype (Figure 2E), indicating the superior capacity of these cells to enter the target organ. However, there were no significant differences between the proportions of either IL-17^−^IFNγ^+^ or total IFNγ^+^ between the sexes at these timepoints.

Granulocyte and macrophage colony-stimulating factor (GM-CSF) is a crucial positive regulator of Th17 cell pathogenicity in both MS and EAE (Becher et al., 2016). As we had observed phenotypic plasticity in male Th17 cells in early disease, we next sought to determine whether male cells produced more of this cytokine. However, no differences in GM-CSF expression between male and female Th17 cells were seen at disease onset or relapse or in either splenic (Supplemental Figure 1A) or CNS (Supplemental Figure 1B) T cells. Similarly, few differences were observed between male and female with respect to TNFα and IL-2 production, though a higher percentage of male splenic Th17 produced TNFα at relapse (Supplemental Figure 1A). We could also rule out an increased conversion of female Th17 to FoxP3^+^ T_reg_, as no differences in the frequency of these cells was observed between males and females (Supplemental Figure 1C).

### Phenotypic plasticity of male Th17 cells underlies a rapid disease course

Our data thus showed important differences between transferred male versus female T cells during early disease timepoints. A substantially greater proportion of splenic male Th17 produced the heterodox cytokine IFNγ as compared to females. This indicated that male Th17 more rapidly adopted a plastic phenotype *in vivo*. However, in the CNS, no differences in IFNγ production were observed between recipients of male or female cells at these timepoints, reflecting the similar kinetics of symptom onset between the two groups. By contrast, we had observed a sex dimorphism at disease endpoint. Mice receiving female 1C6 Th17 universally survived to d70 with disease of moderately severe maximal severity (score 2.6 ± 0.2; Table 1), while there were two possible fates for mice that received male Th17: the majority rapidly attained ethical endpoints (M^rapid^), while a minority survived to d70 (M^d70^) and had disease of comparable severity to those receiving female (F) Th17 (Figure 1C).

We thus examined whether increased Th17 phenotypic plasticity might explain these divergent outcomes, by comparing IL-17 and IFNγ production from T cells isolated from F Th17, M^rapid^ and M^d70^ recipients. In all conditions, as predicted our findings at relapse, the majority of cytokine-producing T cells in both spleen and CNS were IL-17^−^IFNγ^+^. The percentage of splenic Th17 making IFNγ was increased from M^rapid^ mice relative to those from female and M^d70^ (Figure 3A). By contrast, no differences in IL-17 production were observed. Crucially, we saw a large increase in the frequency of IFNγ^+^ T cells in the CNS of M^rapid^ mice, as compared to either F or M^d70^ mice (Figure 3B). Further, CNS-infiltrating Th17 from M^rapid^ made over 12-fold more IFNγ than either female or M^d70^ cells as measured by mean fluorescence intensity (MFI). MFI of GM-CSF, but not of TNFα or IL-2, was also significantly increased from M^rapid^ T cells (Figure 3C). Altogether, our data showed that male sex is an important regulator of Th17 plasticity, and that Th17 production of IFNγ and GM-CSF correlates strongly with disease severity observed in recipients of male Th17.

**Figure 3.**
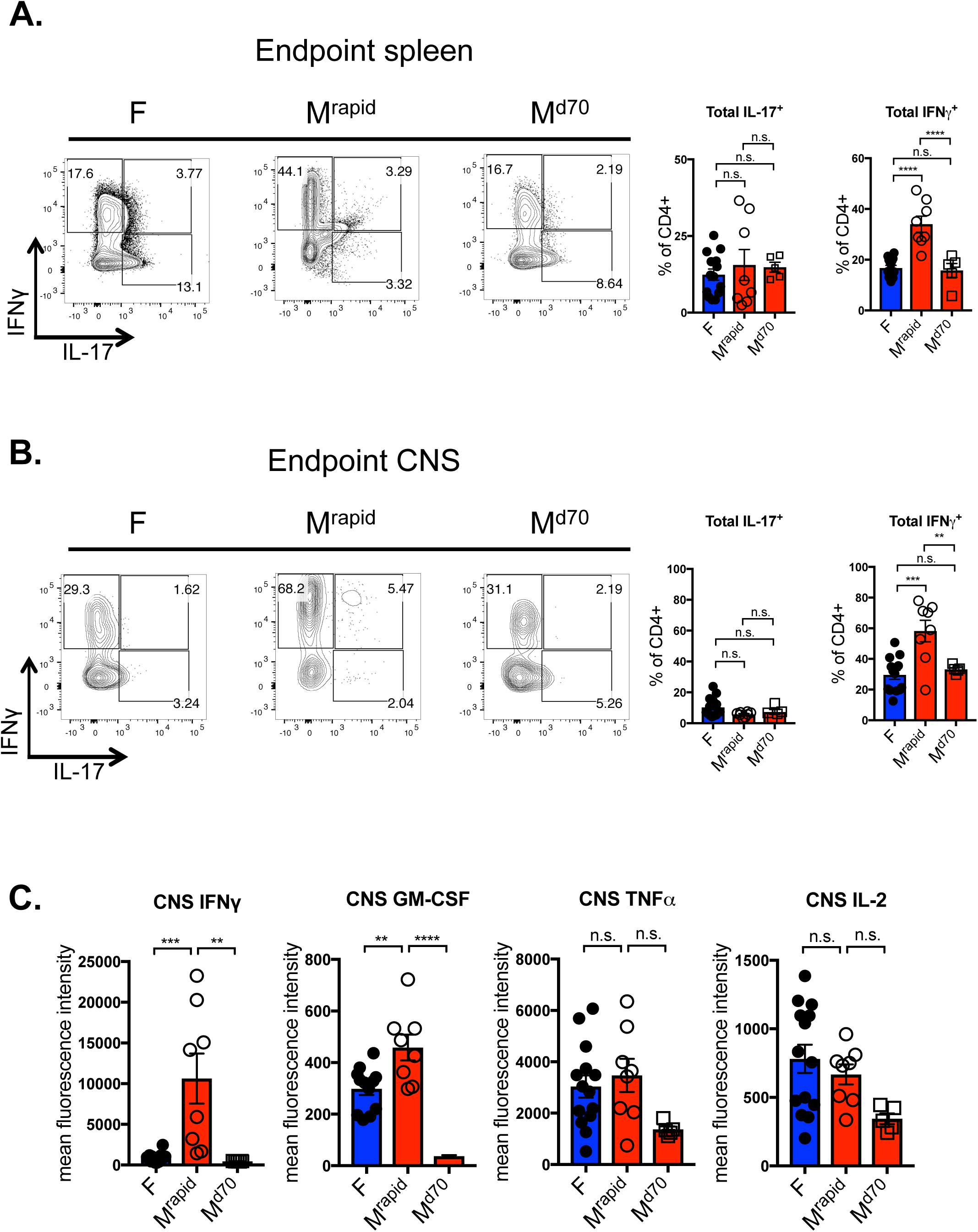
Highly pathogenic male Th17 cells strongly upregulate IFNγ. Recipients of female and male 1C6 Th17 were monitored and sacrificed at experimental endpoints. 8 males (M^rapid^) experienced a rapid and lethal disease course; 5 males (M^late^) survived to d70. All females (n=14) survived to d70. Production of IL-17 and IFNγ from splenic (**A**) and CNS-infiltrating (**B**) CD4^+^ T cells was assessed by intracellular flow cytometry from individual mice. Left plots (**A,B**) are representative of the indicated groups. Gated on live CD4^+^ events with box gates based on fluorescence minus one controls. **C**. Mean fluorescence intensity of the indicated cytokines was measured from CNS samples. **, p<0.01; ***, p<0.001; ****, p<0.0001; Tukey’s multiple comparisons test after one-way ANOVA.

### Male Th1 cells do not induce a rapid disease course

We had found that male Th17 cells induced severe EAE and were highly plastic as characterized by their heightened production of IFNγ relative to female Th17 cells. We therefore wanted to ascertain whether a similar sex dimorphism applies to Th1 cells, which are *bonafide* IFNγ producers and are also potent inducers of EAE in adoptive transfer models (Jäger et al., 2009; Rangachari et al., 2012). We stimulated male and female naïve 1C6 CD4^+^CD62L^hi^ cells with agonistic anti-CD3 and anti-CD28 and under defined Th1 differentiation conditions (Rangachari et al., 2012). After 5 days of culture, the resulting cells were positive for T-bet (Figure 4A), the master Th1 lineage transcription factor (Szabo et al., 2000). We then transferred these Th1 cells to sex-matched NOD.*Scid* recipient mice. Linear regression analysis of the disease curves of host mice suggested that male 1C6 Th1 recipients accumulate symptoms more rapidly than those receiving female Th1 cells (Figure 4B), and maximal disease severity was also higher in mice receiving male Th1 cells (Table 1). Crucially, however, disease severity of male Th1 recipients was significantly lower than that of male Th17 recipients (Table 1), and a greatly reduced proportion of male Th1 recipients experienced a rapid disease course in which mice attained ethical endpoints before 70 days (Table 1). In sum, our data indicated that a sex dimorphism by which male 1C6 Th1 cells worsen the course of EAE relative to female Th1; however, male Th17 cells were required for an extremely severe, accelerated, progressive disease course.

**Figure 4.**
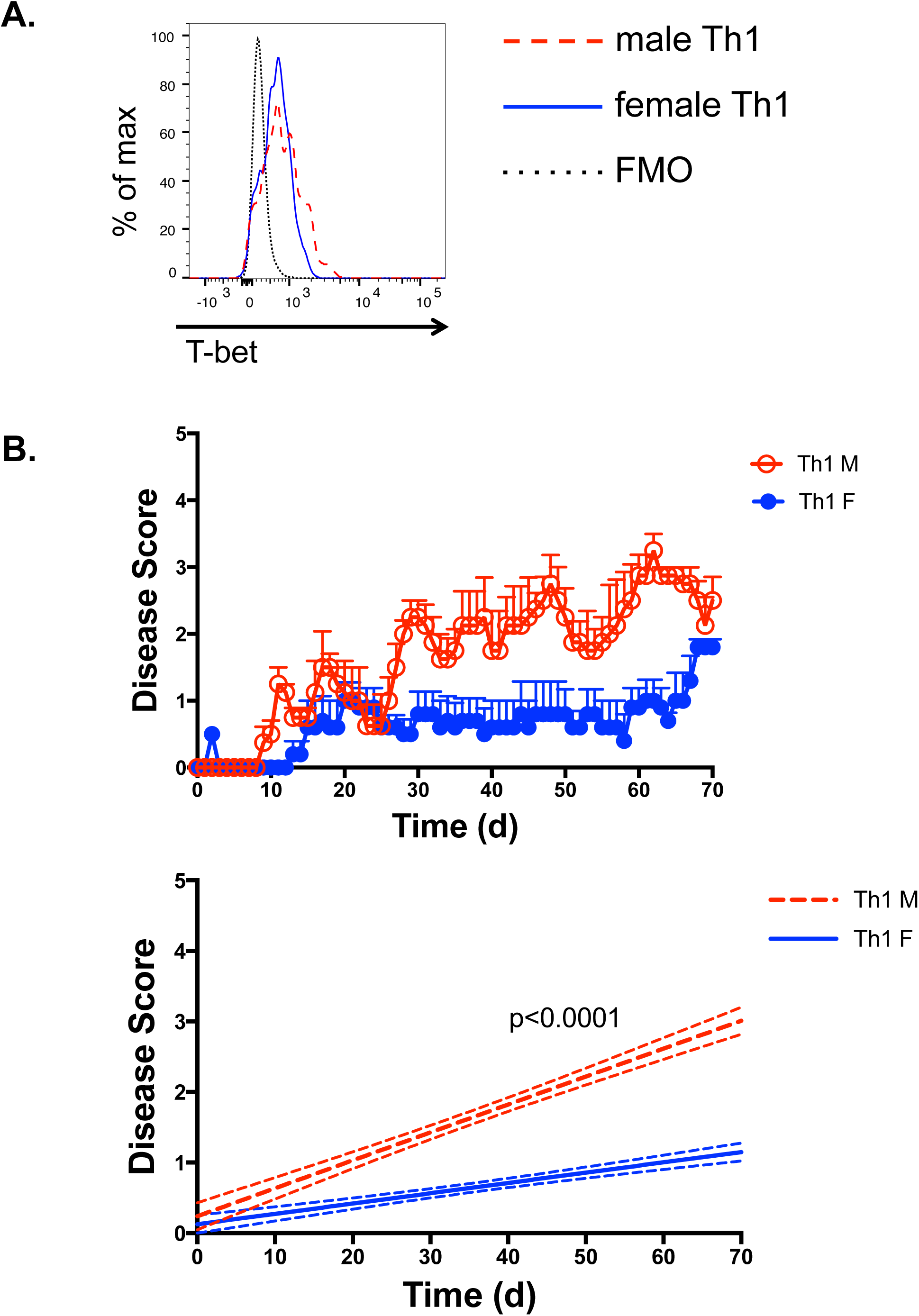
Male Th1 cells do not induce rapid and lethal EAE. Male and female Th1 cells were generated *in vitro* from naïve 1C6 CD4^+^CD62L^hi^ progenitors **A.** T-bet expression was measured by flow cytometry at d5 of culture prior to adoptive transfer. Gated on live CD4^+^ events. Solid blue line, female cells; dashed red line, male cells; dotted black line, FMO (fluorescence minus one) control. Representative of 3 experiments. **B**. Th1 cells were adoptively transferred to sex-matched NOD.Scid male (n=4) or female (n=5) recpients that were monitored for signs of EAE. Day-over-day disease curve of male vs female Th1 recipients (*top*); linear regression analysis of the disease curves (bottom). Representative of 2 transfers.

### Male sex interacts with disease duration, but not severity, in augmenting an ex-Th17 signature

In recent years, there has been intense interest in characterizing diverse Th17 cell fates at the transcriptional level. In an important advance, single cell RNA-seq analyses revealed that Th17 cells can be classified into successive transient states, adopting a Th1-like “ex-Th17” transcriptional profile (“*pre-Th1-like*” → “*Th17/Th1-like*” → “*Th1-like memory*”) as they become pathogenic in C57BL/6J EAE (Gaublomme et al., 2015). We therefore examined whether signature transcripts from these cellular states were upregulated in male Th17 cells. We observed that only 2 of 13 targets – *Il18r1* and *Cxcr6* – were upregulated in splenic male Th17 cells at disease onset (Supplementary Figure 2). Thus, male Th17 did not appear to have initiated a transition towards an ex-Th17 transcriptional profile at the first signs of clinical symptoms, despite a greater frequency of male Th17 being IFNγ^+^ at this timepoint.

We next examined whether the presence of an ex-Th17 transcriptional profile correlated with exacerbated disease outcomes. We compared males that developed rapid, fatal disease (M^rapid^) to females that we sacrificed in parallel (F^early^). In agreement with our previous findings, F^early^ mice did not show signs of severe disease (Figure 5A). We additionally compared males and females that survived to d70 (M^d70^ vs F^d70^). As expected, M^d70^ had disease of comparable severity to F^d70^ (Figure 5B). We then assessed expression of candidate pre-Th1-like (Figure 5C), Th17/Th1-like (Figure 5D) and Th1-like memory (Figure 5E) transcripts from splenic CD4^+^ T cells by qPCR. Of the 13 signature transcripts studied, 5 were upregulated in M^rapid^ Th17 cells relative to F^early^ Th17. Notably, these transcripts included the cell surface receptors *IL18r1* and *Ccr2* (Figure 5C), and *Ifngr1* and *IL2rb* (Figure 5E). To our surprise, M^d70^ Th17 showed strong induction of each stage of the ex-Th17 gene program, despite the fact that these cells transferred EAE of a moderate, non-rapid disease course: eleven transcripts were upregulated in M^d70^ Th17 relative to F^d70^ Th17, with *Il2ra* and *Samsn1* being the sole exceptions. Furthermore, 8 transcripts (*Mina, Fas, Ccr2, Tnfsf11, Ifngr1, Nr4a1, Tigit, Il1r2, Il2ra, Samsn1*) were upregulated in M^d70^ Th17 as compared to M^rapid^ that induced severe disease – in some cases dramatically (Figures 5C-E). Only 3 of these 8 transcripts (*Ifngr1, Il2ra, Samsn1*) were similarly upregulated in F^d70^ versus F^early^, indicating that the correlation between disease duration and pathogenic gene upregulation was principally driven by male sex. Thus, while male sex augments an ex-Th17 gene signature in 1C6 Th17 cells, it does so in relation to disease duration and not to disease severity. Further, the increased production of IFNγ by male Th17 cells that induced rapid, severe, EAE could be uncoupled from the ex-Th17 transcriptional signature.

**Figure 5.**
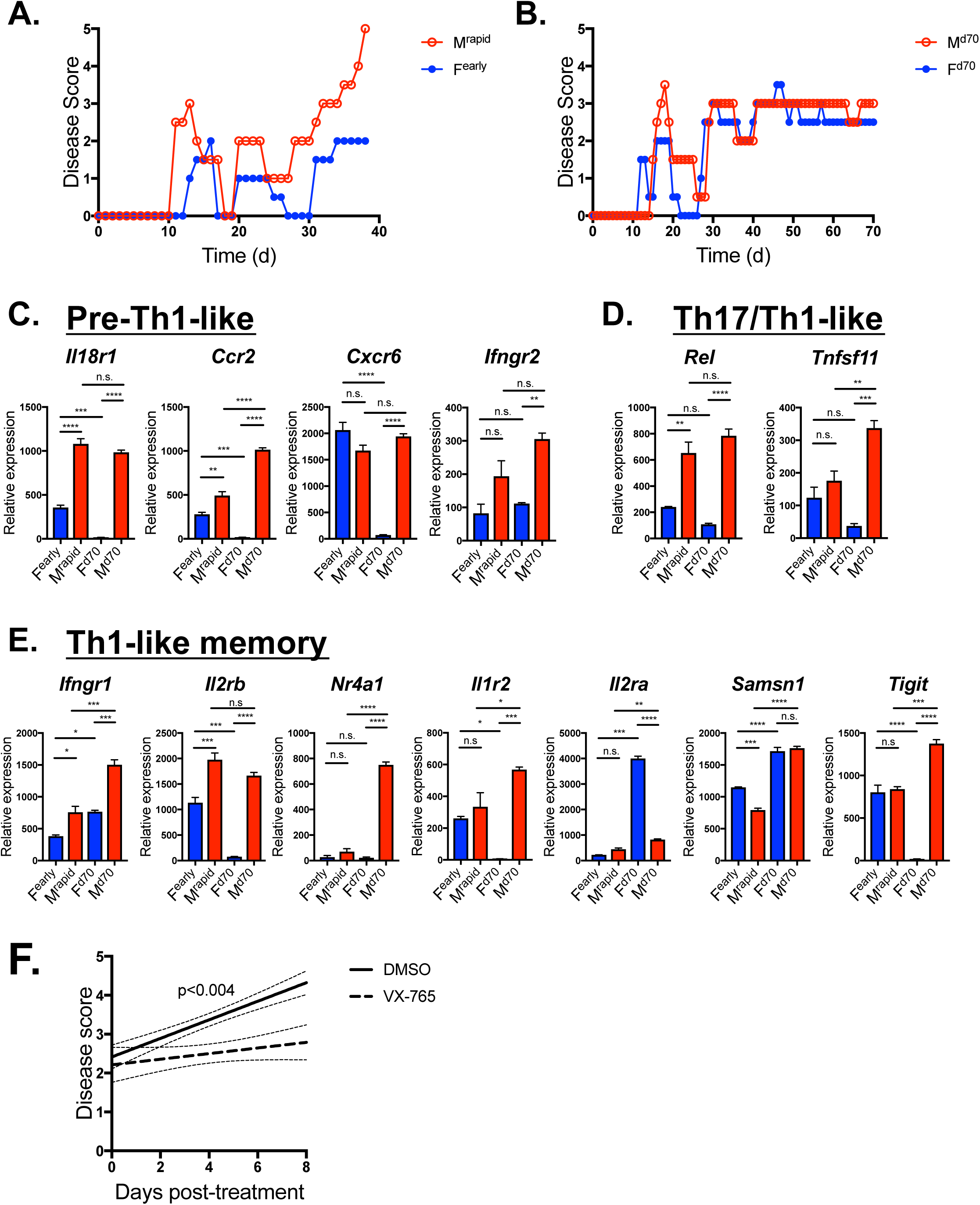
Male sex augments Th17 transcriptional plasticity in a disease duration-dependent manner. Male and female 1C6 Th17 were adoptively transferred to sex-matched NOD.*Scid*. Three males rapidly attained ethical endpoints (M^rapid^). Three females (F^early^) were sacrificed alongside them. The remainder of males (M^d70^, n=5) and females (F^d70^, n=3) were sacrificed at d70. **A**, representative M^rapid^ and F^early^ disease courses. **B**, representative M^d70^ and F^d70^ disease courses. **C, D, E.** cDNA were generated from splenic CD4^+^ T cells and were assessed by qPCR for expression of pre-Th1-like (**C**), Th17/Th1-like (**D**) and Th1-like memory (**E**) transcripts. *, p<0.05; **, p<0.01, ***, p<0.001, ****, p<0.0001, Tukey’s multiple comparisons test after two-way ANOVA. Data represent triplicate values from pooled samples for each condition. Error bars, s.e.m. **F.** Male recipients of male Th17 mice were treated daily for 8 days with inflammasome inhibitor VX-765 (n=6) or DMSO vehicle (n=8) upon attaining score 2. Linear regression analysis of disease curves post-treatment. Error lines represent 95% confidence intervals.

### Inflammasome inhibition modulates the severity of male Th17-induced disease

We next wanted to identify molecules and/or pathways that could be targeted so as to attenuate the severity of disease in male Th17 cell recipients. We noted that the pre-Th1-like signature transcript *Il18r1* was upregulated by more than 2-fold in M^rapid^ Th17 as compared to F^early^ (Figure 5C), and thus might be a possible target. IL-18/IL-18R signaling lies downstream of caspase-1-dependent inflammasome activation, and it was recently shown that the caspase-1 inhibitor VX-765 reduced both inflammasome activation and IL-18 secretion, and improved behavioral outcomes, in the acute C57BL/6J model of EAE (McKenzie et al., 2018). We therefore asked whether inhibiting inflammasome activation *in vivo* could modulate the severe disease induced by male Th17. We transferred male Th17 to male recipients and began treating the mice with VX-765 or vehicle daily once they reached a clinical score of 2 (impaired righting reflex). VX-765 significantly ameliorated the accumulation of disease symptoms in recipient mice with established EAE (Figure 5F), indicating that activation of the inflammasome activity is important for the development of severe disease induced by male 1C6 Th17.

### Chromosome complement exacerbates NOD-EAE in males

Sex hormones can have profound effects on the pathogenesis of EAE (Dunn et al., 2015). It was therefore possible that androgens were responsible for the severe EAE induced by male Th17. We generated Th17 cells from male 1C6 mice that had been castrated (Th17 M^cast^) or sham-operated (Th17 M^sham^), and adoptively transferred them to male NOD.*Scid* mice. Interestingly, we observed that transfer of Th17^cast^ did not reduce disease severity (Figure 6A), indicating that T cell-intrinsic androgens were not pathogenic. We next tested the pathogenicity of Th17 from female 1C6 that had been ovariectomized (Th17 F^ovx^) or sham-operated (Th17 F^sham^), and found that removal of female sex hormones did not worsen EAE in female transfer recipients (Figure 6B). These data showed that differences in pathogenicity between male and female Th17 were not due to sex hormones.

**Figure 6.**
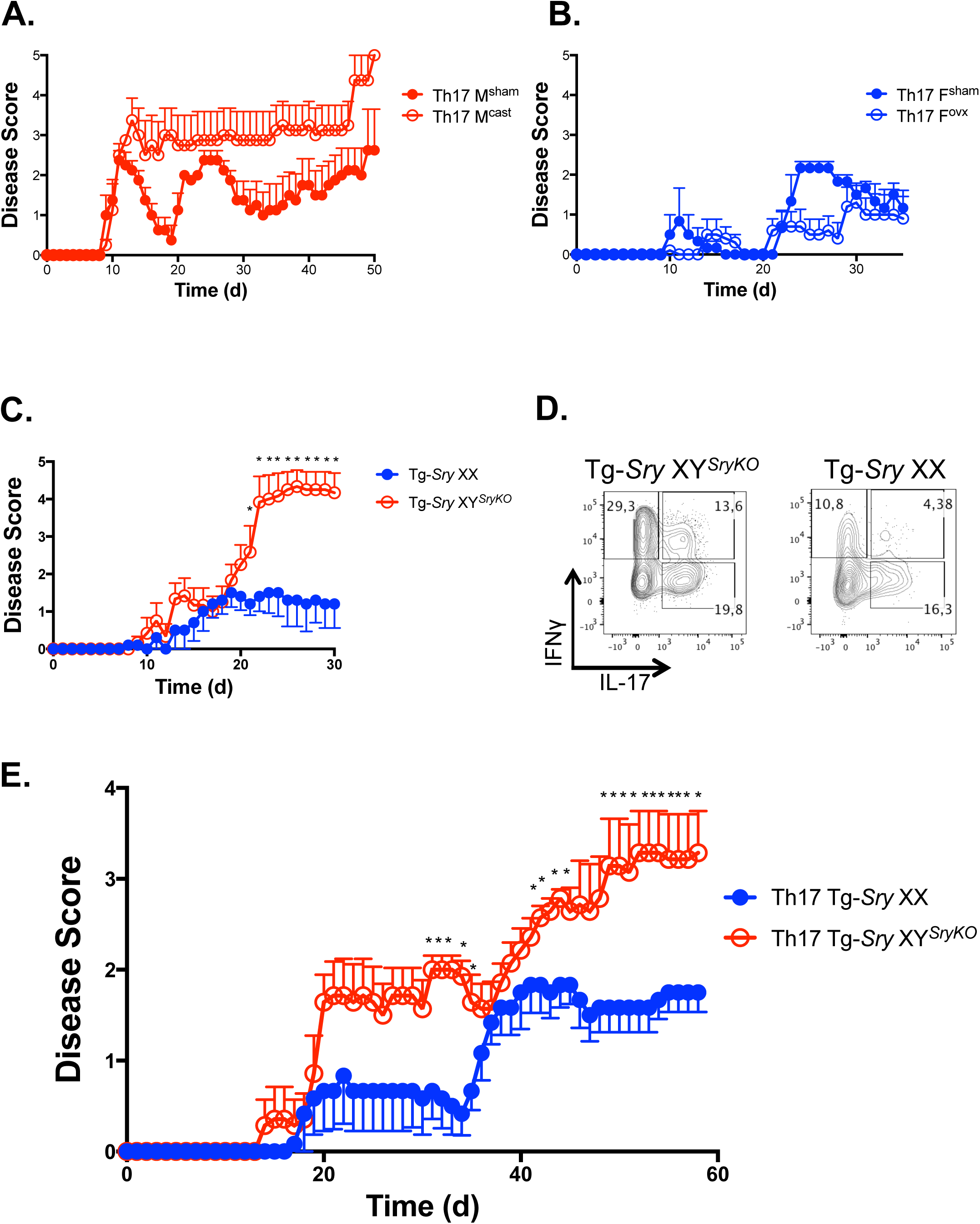
Male sex chromosomal complement aggravates NOD-EAE and a putative X-linked Th17 repressor, *Jarid1c*, is downregulated in highly pathogenic male Th17. **A.** Th17 were generated from castrated male (M^cast^) or sham-operated (M^sham^) 1C6 mice, and were adoptively transferred to male NOD.*Scid* recipients that were monitored for signs of disease. M^cast^, n=4; M^sham^, n=4. **B.** Th17 were generated from ovariectomized female (F^ovx^) or sham-operated (F^sham^) 1C6 mice, and were adoptively transferred to female NOD.*Scid* recipients that were monitored for signs of disease. F^ovx^, n=4; F^sham^, n=4. **C.** Tg-*Sry* XX (n=5) and Tg-*Sry* XY^*SryKO*^ (n=6) were actively immunized with MOG_[35−55]_ and were monitored for signs of EAE. *, p<0.05, Mann-Whitney *U* test. **D**. Co-expression of IL-17 and IFNγ in CNS-infiltrating CD4^+^ from Tg-*Sry* XX or Tg-*Sry* XY^*SryKO*^ (**C)** at experimental endpoint. **E.** Th17 were generated from F1 Tg-*Sry* XX x 1C6 and Tg-*Sry* XY^*SryKO*^ x 1C6 and were transferred (2×10^6^) to male NOD.*Scid* that were monitored for signs of disease. Th17 Tg-*Sry* XX, n=6; Th17 Tg-*Sry* XY^*SryKO*^, n=7.

Sex chromosome complement (XX vs XY) can also play an important role in regulating immune responses (Rubtsov et al., 2010). To assess the role of chromosome complement in the presence of androgen, we exploited the Four Core Genotype (FCG) model (Arnold and Chen, 2009), in which the testes-determining gene *Sry* is knocked out of the Y chromosome (Y^*SryKO*^), but is knocked in to an autosomal chromosome (Tg-*Sry*). We generated F1 NOD x C57BL/6J (B6) Tg-*Sry* XY^*SryKO*^ mice, which produce androgens and are chromosomally male, as well as Tg-*Sry* XX mice, which produce androgens yet are chromosomally female. We then actively immunized these mice with MOG_[35-55]_ and observed their development of EAE, exploiting the prior observation that NOD x B6 F1 mice phenocopy NOD-EAE upon immunization with this encephalitogenic epitope. (Mayo et al., 2014; 2016). We found that Tg-*Sry* XY^*SryKO*^ mice developed disease of dramatically greater severity than Tg-*Sry* XX (Figure 6C). Intriguingly, a substantial frequency of CNS-infiltrating CD4^+^ T cells in Tg-*Sry* XY^*SryKO*^ mice displayed a IL-17^+^IFNγ^+^ phenotype (Figure 6D), which is reflective of Th17 plasticity (Carbajal et al., 2015; Hirota et al., 2011) and pathogenicity (Kebir et al., 2009; Suryani and Sutton, 2007) in active immunization models of EAE.

To directly ascertain whether male sex complement exacerbated the pathogenicity of Th17 cells themselves, we crossed FCG mice to the 1C6 females to generate Tg-*Sry* XY^*SryKO*^ and Tg-*Sry* XX B6 × 1C6 F1 mice. We generated Th17 cells from these lines and adoptively transferred them to male NOD.Scid mice. To mitigate against potential NK cell-mediated host-versus-graft reactivity against B6 antigens (Hummel et al., 2002), we transferred 40% fewer cells (2×10^6^) to recipient NOD.Scid than in our standard protocol. Notwithstanding this lowered dose, Tg-*Sry* XY^*SryKO*^ Th17 induced disease of substantially greater severity that Tg-*Sry* XX Th17 (Figure 6E). Together, these data demonstrated that the male chromosomal complement (XY), and not androgens *per se*, are responsible for promoting the plasticity and pathogenicity of Th17 cells.

### The putative X-linked Th17 repressor Jarid1c is downregulated in highly pathogenic male Th17 and in CD4^+^ T cells of men living with MS

Mammalian female cells inactivate one of their two X-chromosomes, thus ensuring equivalent dosage of X-linked genes between the sexes. However, this process is far from total (Arnold et al., 2016) and numerous genes on the silenced X chromosome continue to be transcribed in a tissue-specific manner (Berletch et al., 2015). Thus, differential dosage of X-linked genes may help explain differences in immune function in females versus males. As our data indicated that male sex chromosome complement exacerbated Th17 pathogenicity (Figure 6E), we hypothesized that X-linked genes might a) repress Th17 pathogenicity, and b) be downregulated in male Th17 cells. To study this possibility, we analyzed publicly available RNA-seq datasets of lymph node and CNS-infiltrating Th17 at the peak of C57BL6/J EAE (Gaublomme et al., 2015). We considered CNS-infiltrating Th17 cells to be pathogenic, as they had infiltrated the target organ. Conversely, lymph node Th17 were considered non-pathogenic. We identified a set of transcripts on the X chromosome that were downregulated by 2-fold or greater in CNS Th17 (Supplementary Table 1) and thus could be inhibitors of Th17 pathogenicity. We then compared these transcripts to a set of X chromosome genes that were previously shown to escape second-copy silencing in female mouse spleen (Berletch et al., 2015), thereby identifying three X-linked transcripts, *Jarid1c, Utp14a* and *Pbdc1* (Figure 7A) that might explain differences in pathogenicity between male and female Th17. Intriguingly, it was recently reported that Jarid1c negatively regulates dendritic cell activation, suggesting that it can suppress immune cell function (Boukhaled et al., 2016).

**Figure 7.**
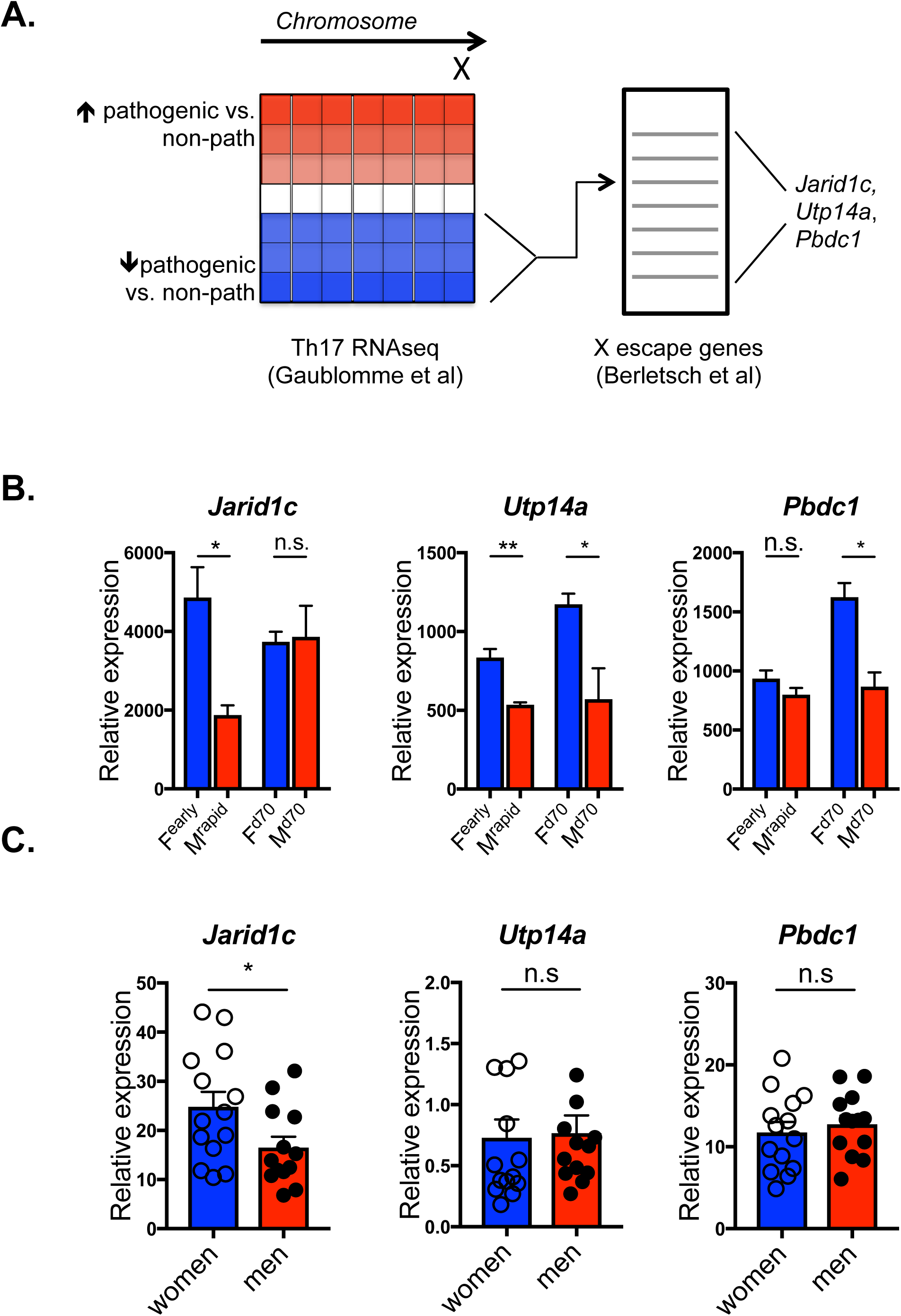
The putative X-linked Th17 repressor Jarid1c is downregulated in highly pathogenic male Th17 and in CD4^+^ T cells of men living with MS. **A**. Schematic of the bioinformatics approach used to identify potential repressors of pathogenic Th17 that are expressed on the X chromosome and that escape second-copy silencing. Jarid1c (downregulated 3.2-fold in CNS-infiltrating Th17), Utp14a (7.7 fold) and Pbdc1 (2.43-fold) were identified. **B**. Expression of putative regulators *Jarid1c, Utp14a* and *Pbdc1* were assessed from CNS-infiltrating CD4^+^ from F^early^ versus M^rapid^ or from F^d70^ versus M^d70^ by qPCR. *, p<0.05; **, p<0.01; t-test. Data represent triplicate values from pooled samples (F^early^, n=3; M^rapid^, n=3; F^d70^, n=3; M^d70^, n=5). Error bars s.e.m. **C.** CD4^+^CD3^+^ T cells were purified from peripheral blood mononuclear cells taken from age-matched men (n=12, 51.4 ± 1.5a) and women (n=14, 52.6 ± 2.2) living with MS. Expression of *Jarid1c, Utp14a* and *Pbdc1* were assessed by qPCR. *, p<0.05, t-test. Each symbol represents an individual.

We next asked whether these putative immunomodulatory X-linked candidates were downregulated in pathogenic male Th17 that induce severe disease. We found that *Jarid1c* was specifically downregulated in CNS-infiltrating Th17 cells from M^rapid^ recipients that develop early, severe disease, as compared to CNS-infiltrating cells from F^early^ mice. (Figure 7B). However, there were no differences in *Jarid1c* expression between Th17 cells from M^d70^ and F^d70^ that developed mild disease of comparable severity. *Utp14a* was downregulated in both M^rapid^ and M^d70^ as compared to timepoint-matched female controls, while *Pbdc1* was only downregulated in M^d70^ (Figure 7B).

We next examined whether sex differences could be observed in the expression of X-linked immune regulators in the context of human MS. We therefore examined expression of *Jarid1c, Utp14a* and *Pbdc1* in purified CD4^+^ T cells from age-matched men (51.4 ± 1.5a) and women (52.6 ± 2.2a) living with MS (Supplementary Table 2). We observed lower *Jarid1c* expression in CD4^+^ T cells from men (Figure 7C). No sex differences were observed in the expression of *Utp14a* and *Pbdc1*. Together, our data showed that male sex complement can worsen progressive EAE, and that the X-linked immunoregulatory gene Jarid1c is downregulated both in pathogenic tissue-infiltrating male Th17 as well as in CD4^+^ T cells from men with MS.

## Discussion

As is the case for many autoimmune diseases, the incidence of RRMS is much (up to 3 times) greater in women than in men (Orton et al., 2006). Nevertheless, men are more likely to progress to a higher level of MS disability than women, and male sex is an exacerbating factor for conversion of RR to SP MS (Koch et al., 2010). As NOD mice develop a chronic progressive form of EAE (Farez et al., 2009; Mayo et al., 2014), we developed an adoptive transfer model on this background to examine whether male sex could increase the incidence and severity of progressive disease in a T cell-intrinsic manner.

We find that recipients of male 1C6 Th17 develop EAE of greater severity than their female counterparts, also developing chronically progressive disease at a higher frequency. While the majority of recipients of male Th17 attain experimental endpoints rapidly, a subset of mice displays disease that is of no greater severity than that of recipients of female Th17. This is important because it allowed us to identify factors that drive worsened pathology in our model. Further, our data show that enhanced disease in males is due to T cell-intrinsic effects, as male 1C6 Th17 cells induce EAE of markedly great severity relative to females regardless of whether they are transferred to male or female hosts. At first glance, this may seem surprising, given that women display a higher CD4^+^ T cell count than men, and that female cells proliferate more robustly upon antigenic stimulation. On the other hand, multiple lines of evidence in mouse (Jane-wit et al., 2008; Li et al., 2013; Zhang et al., 2012) and man (Gracey et al., 2016; Yi et al., 2009; Zhang et al., 2012) indicate that male sex can promote Th17 responses in the context of autoimmunity. Notably, we found that IFNγ production was considerably greater in peripheral male 1C6 Th17 cells at all timepoints studied and crucially, CNS-infiltrating male Th17 cells were highly positive for IFNγ and GM-CSF in mice that develop severe endstage disease. Thus, we demonstrate, for the first time, a role for male sex in augmenting Th17 phenotypic plasticity, which is itself an important determinant of disease severity in autoimmunity (Abromson-Leeman et al., 2009; Carbajal et al., 2015; Y. K. Lee et al., 2009; Suryani and Sutton, 2007). It is worth noting that while female 1C6 Th17 produced less IFNγ *in vivo* than their male counterparts, they also turned off IL-17 expression by the first relapse. This may be characteristic of T cells on the NOD background, as NOD-strain pancreatic Ag-specific Th17 cells are prone to converting to an IL-17^−^IFNγ^+^ phenotype upon transfer to NOD.*Scid*, and blocking IFNγ could rescue Th17-induced diabetes in this model (Bending et al., 2009; Martin-Orozco et al., 2009).

In recent years, increasing attention has been paid to the transcriptional profile of Th17 cells (Gaublomme et al., 2015; Yosef et al., 2013), with the goal of identifying functional gene modules that dictate whether or not they become pathogenic in the context of tissue inflammation. We found that ex-Th17 transcriptional markers were increased in adoptive transferred male Th17 cells, supporting the view that male sex enhances Th17 plasticity. Interestingly, while these markers were initially described as correlating with increased pathogenicity in the context of EAE (Gaublomme et al., 2015), we found that while some were indeed upregulated in male Th17 that transferred severe disease, they were more strongly associated with male Th17 that induced relatively milder disease in recipients that survived to d70. A potential explanation for this discrepancy is that these targets were initially identified using an IL-17-GFP reporter mouse strain in which Th17 cells were actively generating IL-17 at the time of harvest (Gaublomme et al., 2015). We, on the other hand, observe that adoptively transferred 1C6 Th17 lose IL-17 expression relatively early on in disease. Thus, it is possible that this transcriptional signature may correlate with plasticity, but not necessarily pathogenicity, in Th17 cells that have lost IL-17 expression entirely. Importantly, IFNγ (Abromson-Leeman et al., 2009; Carbajal et al., 2015; Hirota et al., 2011; Suryani and Sutton, 2007) and GM-CSF (Becher et al., 2016), which are functional markers of pathogenic Th17 cells, are strongly upregulated in male Th17 that transfer severe disease, suggesting that pathogenicity of these cells may be uncoupled from transcriptionally-defined plasticity in our model. Another important question is why a proportion (∼30%) of male Th17 recipients develop only mild disease. Overall, splenic CD4^+^ from male Th17 recipients showed increased IFNγ at onset and relapse; it is possible that differential expression of this cytokine, or of specific plasticity transcripts, at early timepoints might predict disease outcomes. However, the terminal nature of these analyses makes it challenging to address this directly. Further studies will be required to address this issue, perhaps using IL-17 fate-mapping mice (Hirota et al., 2011).

The elegant FCG mouse model permits one to separately consider the contributions of sex hormones and sex chromosomes to biological processes (Arnold and Chen, 2009). This is important because, in contrast to sex hormones, the role played by sex chromosomes is often less well understood (Palaszynski et al., 2005). In agreement with previous studies of active immunization EAE (Bebo et al., 1998; Palaszynski et al., 2004; Trooster et al., 1996), we found that castration of male 1C6 mice did not reduce the pathogenicity of Th17 derived from these animals. Thus, we exploited the FCG model to generate Tg-*Sry* XX and Tg-*Sry* XY^*SryKO*^ (NOD x B6 F1) 1C6 mice, and found that Tg-*Sry* XY^*SryKO*^ 1C6 Th17 transferred far more severe EAE to NOD.Scid recipients. Thus, we show for the first time that T-cell-intrinsic, sex chromosome-encoded, factors regulate sex-specific differences in Th17 cell plasticity and pathogenicity. It must be noted that the contribution of male sex chromosomes is highly strain-dependent, as the XY complement is actually immunoprotective in both active immunization and adoptive transfer EAE on the SJL/J strain, while sex chromosome complement does not affect C57BL6/J EAE (Smith-Bouvier et al., 2008). The interplay of sex chromosomes and hormones is therefore complex and strain-specific, with different genetic backgrounds revealing distinct pathogenic or regulatory mechanisms in the pathogenesis of EAE. In the case of the NOD background, sex chromosome factors may interact genetically with *Idd* disease susceptibility loci (Wicker et al., 1995). In further studies, it will be instructive to examine the contribution of these loci to worsened Th17-mediated disease in males.

In an elegant recent study, it was shown that when transferred to syngeneic SJL/J hosts, splenocytes from *in vivo*-primed male, MOG-specific, transgenic TCR^1640^ mice induced progressive EAE while female cells transferred RR-EAE (Dhaeze et al., 2019). In contrast to our findings, however, male splenocytes in this model did not appear to transfer disease of increased maximal severity relative to female cells. While IL-17 and IFNγ production were not directly compared between transferred male and female TCR^1640^ T cells, it was observed that the frequency of IL-17^+^IFNγ ^−^ cells diminished in males as disease progressed while the frequency of IL-17^−^ IFNγ^+^ increased. An important difference between this study and our own was that prior to transfer, Dhaeze and colleagues stimulated splenocytes with IL-12 as well as IL-23 so as to reactivate both Th1 and Th17 cells. Thus, it probable that the IL-17^−^IFNγ^+^ cells that they identified at chronic disease in males comprised a mixture of both *bonafide* Th1 cells as well as ex-Th17 cells. By contrast, we specifically generated Th17 cells that were highly Rorγt-positive prior to adoptive transfer, and male 1C6 Th17 generated under these conditions induced an accelerated disease course of greater severity than that invoked by male Th1. Our data thus uncover a crucial role for the Th17 compartment specifically in mediating sex differences in the severity of progressive CNS autoimmunity. Further, we pinpoint a role for sex chromosome factors in regulating this dimorphism.

Using publicly available transcriptomics data and a bioinformatics-based approach, we have identified 3 genes on the X chromosome, *Jarid1c, Utp14a* and *Pbdc1*, that escape second-copy silencing in the immune compartment (Berletch et al., 2015) and are additionally downregulated in pathogenic T cells. Further, we find that expression of one of these genes, Jarid1c, is profoundly downregulated in male Th17 cells taken from mice that experience lethal disease. Jarid1c is a histone H3K4 demethylase that promotes gene transcription when binding to enhancer regions, yet represses transcription when binding to core promoter sequences (Outchkourov et al., 2013). In a recent advance, it was shown that Jarid1c has an immunomodulatory function by enforcing a state of dendritic cell quiescence in collaboration with the transcriptional repressor Pcgf6 (Boukhaled et al., 2016). It will be important in the future to understand whether Jarid1c is a broad repressor of leukocyte activation and whether its expression in dendritic cells is also regulated in a sex-specific manner.

While here we identified potential X-linked regulators of Th17 pathogenicity, it is important to note that Y chromosome loci, such as Yaa, can strongly influence autoimmune disease (Izui et al., 1995). A Y-chromosome element regulates the resistance of young SJL/J males to EAE (Spach et al., 2009). Intriguingly, the Y chromosome can have genome-wide imprinting effects: female mice display the EAE susceptibility of their male sires or grandsires (Teuscher et al., 2006) potentially as a result of copy number variation of Y chromosomal multicopy genes (Case et al., 2015). It would interesting in the future to pinpoint a potential role for the Y chromosome in regulating disease severity in our model, perhaps via its genetic interaction with *Idd* loci (Wicker et al., 1995).

In conclusion, we show that male Th17 cells induce severe progressive CNS autoimmunity while adopting a plastic phenotype *in vivo*. Male Th17 pathogenesis is driven by sex chromosomal complement rather than by androgens. These findings represent an important advance in our understanding of how biological sex acts as a determinant of disease severity in CNS autoimmunity.

## Materials and Methods

### Ethics statement

All experimental protocols and breedings were approved by the Animal Protection Committee of the Centre de recherche du CHU de Québec - Université Laval (2017-037-2 and 2017-090-2; both to *M.R.*). All experiments involving human participation were approved by the Newfoundland Health Research Ethics Board with informed consent (to *C.S.M.*). MS patients were recruited through the Health Research Innovation Team in Multiple Sclerosis (HITMS) at Memorial University of Newfoundland, St. John’s, Canada.

### Reagents and cytokines

Flow cytometry monoclonal Abs (mAbs) against mouse antigens were obtained from eBioscience (CD4, clone RM4-5; CD3, clone 145-2C11; CD62L; clone MEL-14; IFNγ, clone XMG1.2; TNFα, clone MP6-XT22; GM-CSF, clone MP1-22E9; FoxP3, clone FJK-16S), Biolegend (IL-17A, clone TC11-18H10.1; T-bet, clone 4B10), BD (IL-2, clone JES6-5H4) or Invitrogen (RORγt, clone AFKJS-9). The following mAbs, all obtained from BioXcell, were used for *in vitro* T cell cultures: anti-CD3 (clone 145-2C11), anti-CD28 (clone 37.51), anti-IFNγ (XMG1.2) and anti-IL-4 (clone 11B11). The following recombinant cytokines used for *in vitro* T cell cultures: recombinant human (rh) TGFβ (Miltenyi), recombinant mouse (rm) IL-6 (Miltenyi), rmIL-23 (R&D Biosystems), rmIL-12 (R&D Biosystems), rmIL-2 (Miltenyi). Inflammasome inhibitor VX-765 was obtained from AdooQ Biosciences.

### Animals, EAE induction and monitoring

1C6 mice were a kind gift of Dr. Vijay Kuchroo (Boston, MA). NOD.*Scid* mice were purchased from Jackson Laboratories (JAX) and were bred in our animal facility. CD4^+^ T cells were isolated from 1C6 mice at 8-14 weeks of age. NOD.*Scid* mice were used between 9-12 weeks of age. Where required, 1C6 were gonadectamized (castration/ovariectomy) or sham-operated at 3 weeks of age. For adoptive transfer of effector T cells, 5×10^6^ Th17 or Th1 1C6 T cells were injected i.p., in cold PBS (Cellgro) into NOD.*Scid* recipient mice, which additionally received 200 ng of pertussis toxin (List Biological Labs) i.p. on d0 and d2 post-injection. Four Core Genotype (Tg-*Sry* XY^*SryKO*^) mice on the C57BL/6J (B6) background were purchased from JAX and were bred to NOD or 1C6 females to obtain Tg-*Sry* XX and Tg-*Sry* XY^*SryKO*^ mice. FCG B6 x NOD F1 mice were actively immunized with 200 µg MOG_[35−55]_ in 100 µL of complete Freund’s adjuvant (Difco) supplemented with 500 µg *M. tuberculosis* extract (Difco), and additionally received 200 ng pertussis toxin (List Biological Labs) on d0 and d2. Th1 and Th17 cells generated from FCG B6 × 1C6 mice were adoptively transferred to NOD.*Scid* at 2.5 × 10^6^ cells/recipient. All EAE mice were weighed and assessed for disease symptoms daily for up to 70 days, using an established semi-quantitative scale that we have used previously (Boivin et al., 2015): 0, no disease; 1, decreased tail tone; 2, hind limb weakness or partial paralysis; 3, complete hindlimb paralysis; 4, front and hind limb paralysis; 5, moribund or dead. Moribund mice were euthanized with 24 hours if they did not show improvement, and were considered score 5 from that point until the end of the experiment. “Disease onset” is defined as being within 24 hours of the first signs of clinical symptoms. “Relapse” is defined as an increase in clinical score of ≥1 that lasts ≥2 days; “remission” as a reduction of ≥1 that lasts ≥2 days (McCarthy et al., 2012). “Rapid course” is defined as attaining ethical endpoints prior to d70. A mouse was determined to have undergone chronic worsening disease if it a) increased in severity of ≥1 over a 10-day period, *and* b) at no point during the 10-day period had a day-over-day decrease of >0.5 in score, *and* c) never remitted after the 10-day period *and* d) passed a threshold of score 2.5 either during or after the 10-day window. Disease was considered “relapsing/chronic” if it featured at least one relapse, in addition to the initial onset, followed by progressive disease. For inflammasome inhibition experiments, male mice were adoptively transferred with male Th17. Mice were alternately placed in treatment (VX-765; 50 mg/kg) or vehicle (DMSO) groups upon attaining score 2. Treatment was maintained for 8 consecutive days.

### T cell purification and culture

Mononuclear cell preparations were obtained from the spleens and lymph nodes (LN) of donor mice. CD4^+^ T cells were enriched using magnetic, anti-CD4 coated beads (Miltenyi) and were subsequently incubated with fluorochrome-labeled anti-CD62L. Naïve CD4^+^CD62L^hi^ T cells were purified using a FACSAria (BD) high-speed cell sorter and were cultured in DMEM (Life Technologies) with 10% fetal calf serum (Corning) and supplemented as described (Sabatos et al., 2003). Cells were stimulated with 2 µg mL^−1^ each of plate-bound anti-CD3 and anti-CD28, under Th17 (3 ng mL^−1^ rhTGFβ + 20 ng mL^−1^ rmIL-6 + 20 µg mL^−1^ anti-IFNγ) or Th1 (10 ng mL^−1^ rmIL-12 + 10 µg mL^−1^ anti-IL-4) conditions. After 2 days, cells were re-plated in the absence of agonistic anti-CD3/CD28 but in the presence of 20 ng mL^−1^ rmIL-23 (for Th17) or 10 ng mL^−1^ rmIL-2 (for Th1) for an additional 3 days. Cells were collected and washed twice in cold PBS prior to injection.

### Histopathology

Mice were euthanized and perfused first with cold PBS and then with 4% paraformaldehyde (PFA) through the left cardiac ventricle. Brains, optic nerves and spinal cords were dissected from the skull and spinal column, respectively. Tissues were stored for 24 hours in 4% PFA at 4°C and then for a minimum of 48 hours in PBS before being embedded in paraffin. Five µm thick sections of the brain, optic nerve and the spinal cord were stained with hematoxylin & eosin (H&E), Luxol fast blue for myelin or Bielschowsky’s silver impregnation for neurons and axons (Bauer and Lassmann, 2016). The inflammatory infiltrate, and area of demyelination, per section was determined from 10 to 12 complete sections per cervical, thoracic and lumbar spinal cord and from complete cross sections of the optic nerve. For inflammation we counted the average number of perivascular inflammatory infiltrates per spinal cord cross sections. The area of demyelination was determined using a morphometric grid (Oji et al., 2016) at an objective lens magnification of 10x in sections stained with luxol fast blue by counting the number of grid points located of the area of myelin loss. The mean value per animal was determined by taking the average values for all sections from a given animal.

### CNS mononuclear cell isolation

Mice were euthanized and perfused with cold PBS administered through the left cardiac ventricle. Brains and spinal cords were dissected from the skull and spinal column, respectively. CNS tissue were homogenized using a PTFE Tissue Grinder (VWR) and were incubated for 30 minutes at 37°C in homogenization solution (HBSS containing 4 ng mL_−1_ liberase and 25 ng mL_−1_ DNase). Homogenate were filtered through a 70 µm cell strainer, resuspended in 35% Percoll (GE Healthcare) and centrifuged. Mononuclear cells were collected, washed and prepared for flow cytometric analysis. For qPCR experiments, cells were stained with fluorochrome-labeled Abs to CD4, CD3 as well as Live/Dead indicator, and live CD4^+^CD3^+^ cells were purified by high-speed cell sorting as described above.

### Flow cytometry

Staining for cell surface markers was carried out for 30 minutes at 4°C. Prior to incubation with mAbs against cell surface markers, cells were incubated for 10 minutes in the presence of Fc Block (BD Biosciences) to prevent non-specific Ab binding to cells. For analysis of intracellular cytokine expression, cells were first cultured for 4 hours in the presence of 50 ng mL^−1^ phorbol 12-myristate 13-acetate (Sigma-Aldrich), 1 µM ionomycin (Sigma-Aldrich) and GolgiStop (1 µL per mL culture; BD Biosciences). Cells were subsequently incubated with Fc Block (Biolegend) and fluorescent cell surface marker Abs and were then fixed and permeabilized using Fixation and Perm/Wash buffers (eBioscience). They were then stained with fluorescent Abs against intracellular markers. True-Nuclear Transcription Buffer Set (Biolegend) was used for RORγt and T-bet staining. FoxP3 Fix/Perm and Perm buffers (Biolegend) were used for FoxP3 staining. Flow cytometry data were collected using an LSRII flow cytometer (BD Biosciences) and were analyzed using FlowJo software (Treestar). Dead cells were excluded from analysis on the basis of positivity for Fixable Viability Dye (eBioscience). Gates were set on the basis of fluorescence minus one controls.

### Human CD4^+^ T cell isolation, RNA extraction, and gene analysis

Cryopreserved human PBMCs were thawed, quickly washed in sorting buffer (1% BSA in PBS, 1mM EDTA, 10nM HEPES), and counted. Cells were then stained with anti-CD3 and anti-CD4 antibodies (BD Biosciences), in addition to Aqua Fluorescence Reactive Dye (ThermoFisher) according to manufacturer’s protocol. Following a wash, stained CD3^+^CD4^+^ cells were sorted using FACS (Astrios, BD Biosciences) and immediately placed in Trizol LS (ThermoFisher). RNA extraction and DNAse treatment was performed according to manufacturer’s instruction using a RNeasy Micro kit (Qiagen). Purified RNA (200ng) was reverse-transcribed to cDNA (M-MLV RT kit; ThermoFisher).

### Quantitative PCR

RNA were isolated from purified CD4^+^ T cells using Direct-zol RNA Purification Kit (Zymo Research) and cDNA were generated using iScript Reverse Transcriptase (Bio-Rad). qPCR was carried out on a Lightcycler 480 II (Roche) using Taqman validated FAM-labeled probes and Gene Expression Master Mix (Applied Biosystems). Samples were run in triplicate. *Rps18* was used as a housekeeping control for mouse studies (Axtner and Sommer, 2009; Cooley et al., 2015). For analysis of human CD4+ T cells, qPCR was performed on reverse-transcribed cDNA using Taqman-validated probes for Jarid1c and the housekeeping genes 18S and GAPDH, on a Viia7 Real-Time PCR System (ThermoFisher). The mean of 18S and GAPDH values was used a reference for human studies.

### Bioinformatics analysis

We retrieved transcriptomic data from GEO project GSE75105 (Gaublomme et al., 2015). The six Paired-end RNAseq fastq datasets were trimmed using trimmomatic v0.36 using this command:

*java -jar -Xmx14G trimmomatic-0.36.jar PE -threads 8 -phred33 sample_R1.fastq.gz sample_R2.fastq.gz trimmed/fastq_R1.fastq.gz trimmed/sample_R1_unpaired.fast trimmed/sample_R2.fastq.gz trimmed/sample_R2_unpaired.fastq.gz*

*ILLUMINACLIP:adapters.fa:2:30:10 TRAILING:30*

*SLIDINGWINDOW:4:20 MINLEN:30*.

Then, trimmed fastq files were mapped on *Mus musculus* (taxid: 10090) Ensembl genome version 92 (Mus_musculus.GRCm38.cdna.standard_chr.fa.gz.idx) using Kallisto 0.44.0. Finally, differential expression between groups CNS and LN were performed using DESeq2 v1.18.1

### Statistical analysis

Two-tailed comparisons were made in all cases. For EAE data, *t*-test was used to analyze mean day of onset and mean number of relapses. Mann-Whitney *U* test was used to analyze mean maximal severity and severity on individual days. ANOVA and Dunn’s multiple comparisons test was used when three or more groups were compared. Fisher’s exact test was used to analyze disease incidence, mortality incidence and progressive phase incidence. Bonferroni’s correction was applied when three groups were compared. Slopes of the disease onset curve were obtained and analyzed by performing linear regression (Boivin et al., 2015). Flow cytometric (percentages of marker-positive cells) and qPCR data were analyzed by Student’s *t*-test. ANOVA and Tukey’s multiple comparisons test was used when three or more groups were compared. Histopathological quantification data were analyzed by Mann-Whitney *U* test. All statistical analyses were performed using Prism (GraphPad), with the exception of Fisher’s exact test (QuickCalc online tool, https://www.graphpad.com/quickcalcs/contingency1.cfm; GraphPad). Error bars represent standard error (s.e.m.).

## Acknowledgements

We thank Vijay K. Kuchroo for providing us with 1C6 mice; Mohammad Balood, Alexandre Brunet and Julie-Christine Lévesque for technical assistance; and Julie Savage and Youjin Lee for critical review of the manuscript. P.M.I.A.D is supported by a Doctoral Studentship from the Multiple Sclerosis Society of Canada (MSSOC). M.R. is supported by a *«Junior-2 chercheurs-boursiers»* salary support award from the Fonds de Recherche de Québec – Santé (FRQS). The work was funded by an Operating Grant (#3036) from the MSSOC as well as by a Catalyst Grant on Sex and Gender as Variables in Biomedical Research from the Canadian Institutes of Health Research (CIHR), both to M.R.

---

**Supplemental Figure 1.**
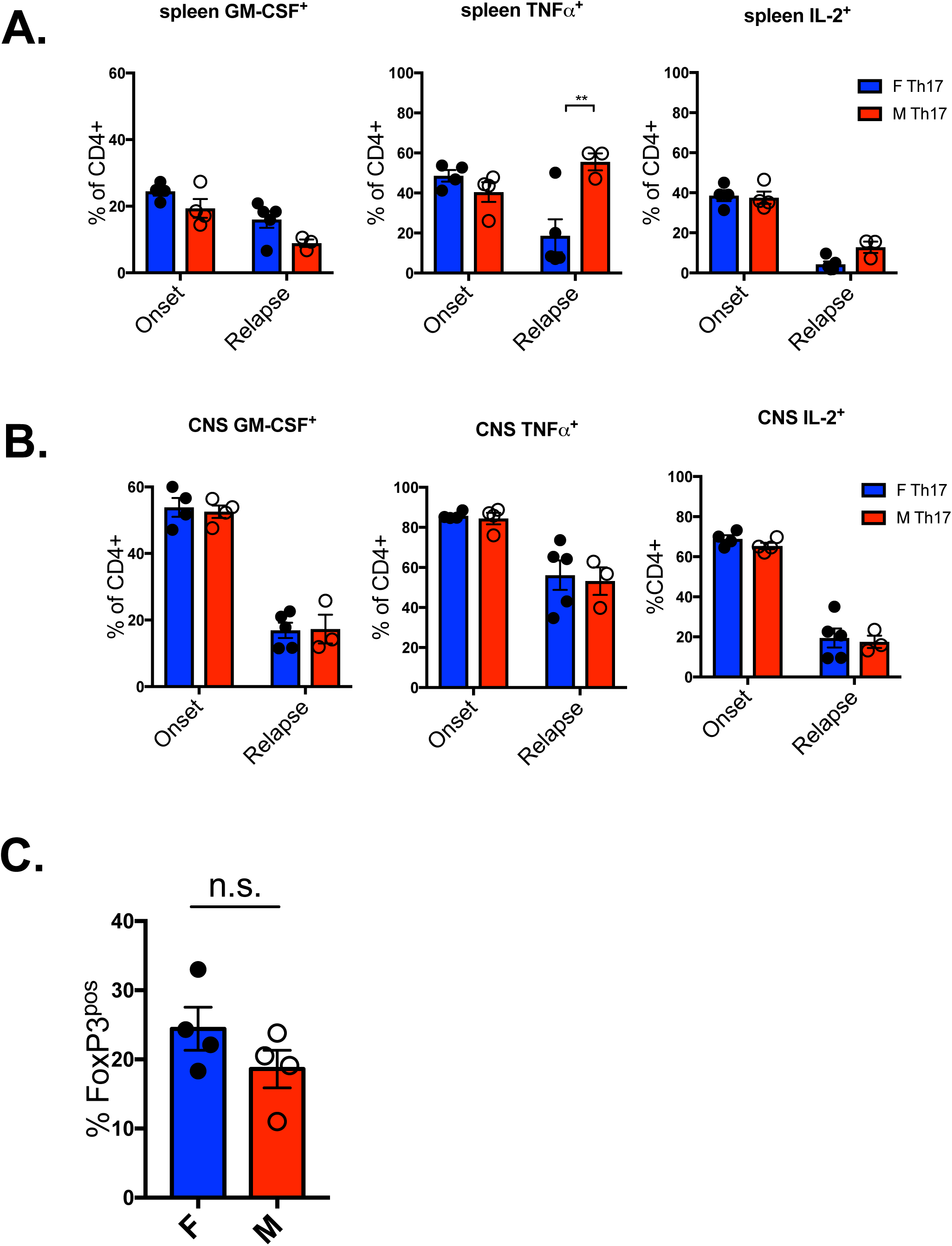
Female recipients of female Th17 or male recipients of male Th17 were sacrificed at disease onset (F, n=4; M, n=4) and relapse (F, n=4, M, n=3). Production of GM-CSF, TNFα and IL-2 were assessed from splenic (A) or CNS-infiltrating (B) CD4^+^ T cells. **, p<0.01, t-test. Gated on live CD4+ events. C. FoxP3^+^ CD4^+^ T cells were assessed from the spleens of female vs. male recipients at onset. Gated on live CD4^+^ events.

**Supplemental Figure 2.**
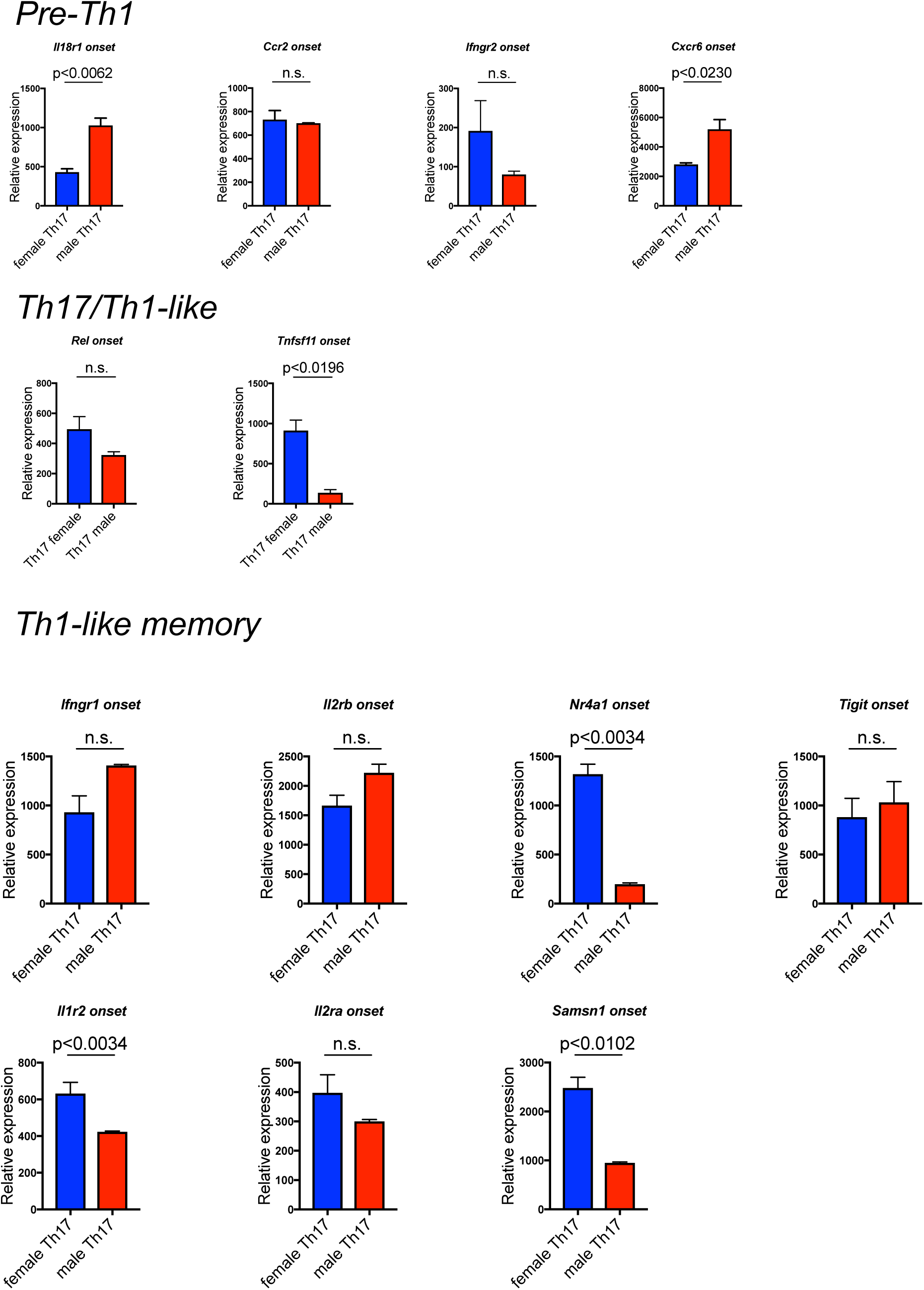
Female recipients of female Th17 or male recipients of male Th17 were sacrificed at disease onset and expression of pre-Th1-like, Th17/Th1-like and Th1-like memory transcripts was assessed from splenic CD4^+^ T cells by qPCR. Data pooled from 3 female and 3 male mice. *t*-test was applied.

**Supplemental Table 1. X chromosome transcripts that are downregulated by 2-fold or greater in pathogenic versus non-pathogenic Th17.** RNA-seq data from CNS-infiltrating (pathogenic) versus LN (non-pathogenic) Th17 cells(Gaublomme et al., 2015) were analyzed to identify transcripts on the X chromosome that were downregulated by 2-fold or greater in pathogenic cells. Data presented by ascending order of expression in pathogenic cells. Transcripts of interest in this study (*Utp14a, Kdm5c, Pbdc1*) that escape X-linked silencing in spleen (Berletch et al., 2015) are bolded and italicized in the list. *Kdm5c* is the gene name of Jarid1c.

**Supplemental Table 2. Characteristics of MS patients analyzed.** M, male; F, female. RR, relapsing/remitting; SP, secondary progressive; PP, primary progressive.

