## Supplementary figures and images for "Male sex chromosomal complement exacerbates the pathogenicity of Th17 cells in a chronic model of CNS autoimmunity"

### Supplemental Figure 1

#### Slide 1
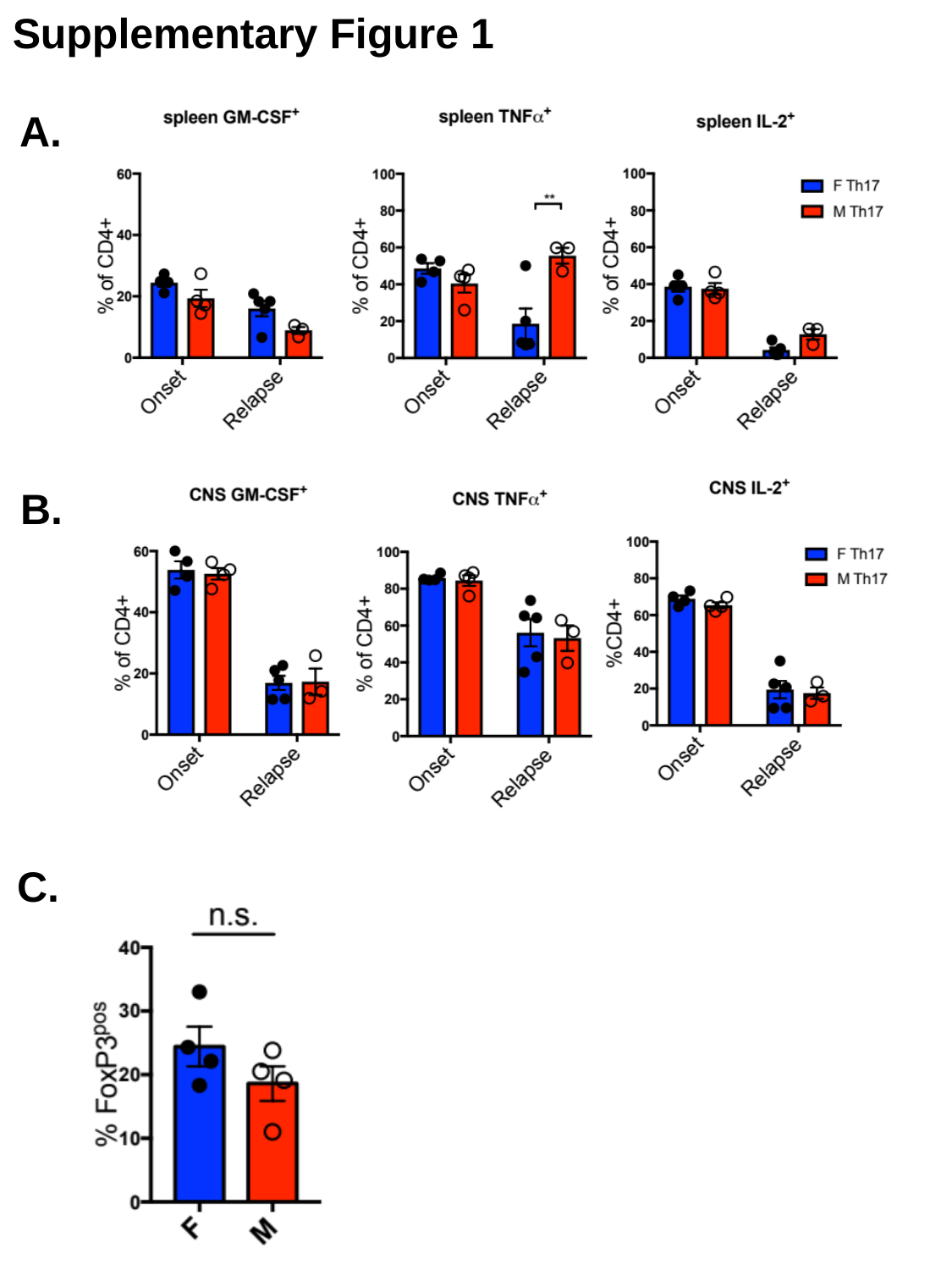

Supplementary Figure 1
A.
B.
C.

### Supplemental Figure 2

#### Slide 1
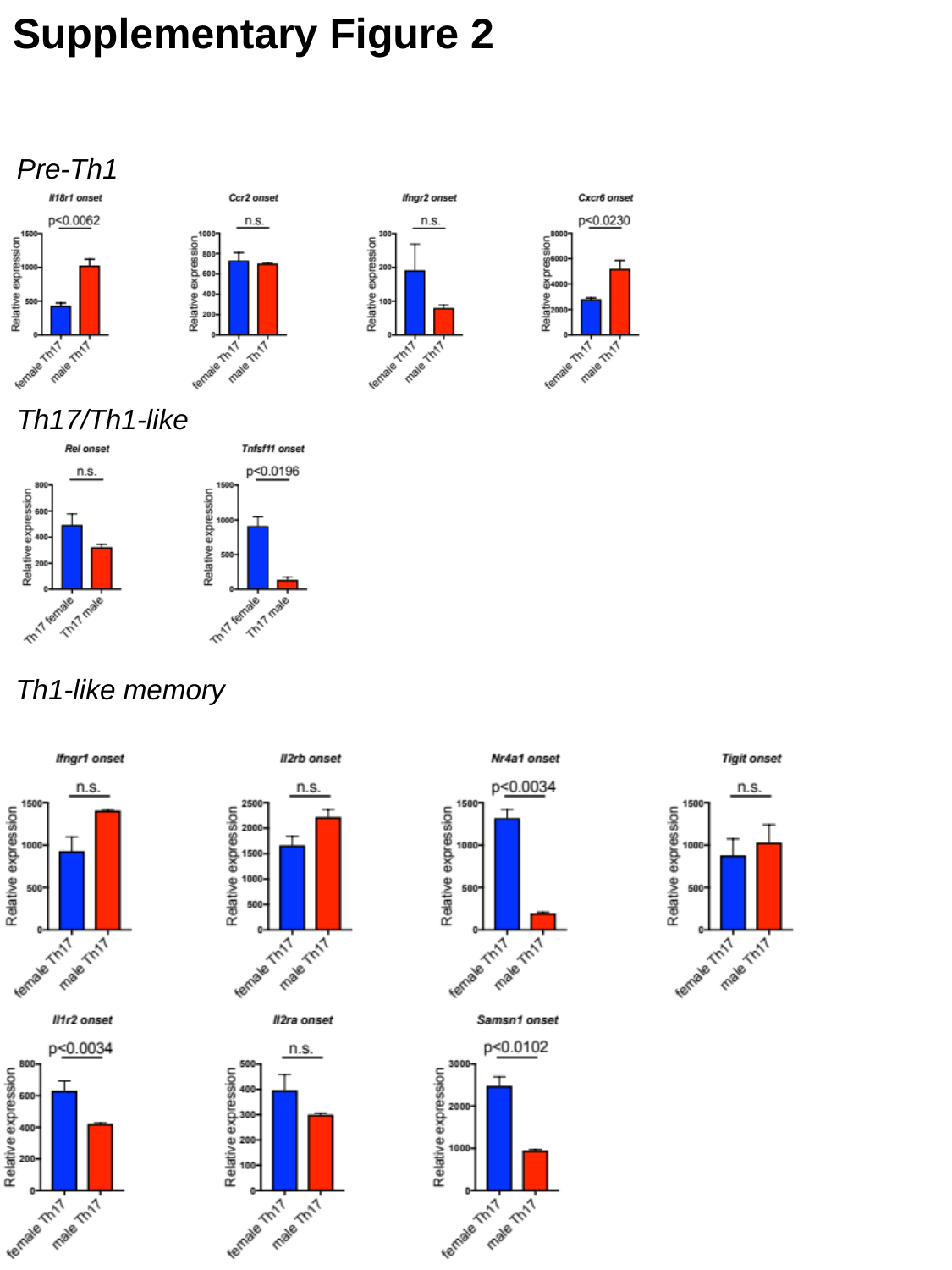

Supplementary Figure 2
Pre-Th1
Th17/Th1-like
Th1-like memory
