## Supplemental Table 2 for "Male sex chromosomal complement exacerbates the pathogenicity of Th17 cells in a chronic model of CNS autoimmunity"

| **Sex** | **Age** | **Type of MS** |
| --- | --- | --- |
| M | 52 | RR |
| M | 41 | SP |
| M | 55 | PP |
| M | 59 | RR |
| M | 52 | RR |
| M | 47 | RR |
| M | 45 | RR |
| M | 62 | RR |
| M | 53 | RR |
| M | 42 | PP |
| M | 51 | SP |
| M | 54 | PP |
| M | 56 | RR |
| F | 50 | RR |
| F | 44 | SP |
| F | 55 | SP |
| F | 61 | RR |
| F | 31 | RR |
| F | 55 | RR |
| F | 45 | RR |
| F | 55 | SP |
| F | 63 | RR |
| F | 52 | RR |
| F | 55 | SP |
| F | 54 | SP |
| F | 59 | PP |
| F | 57 | RR |

**Supplemental Table 2. Characteristics of patients analyzed in Figure 7.** M, male; F, female. RR, relapsing/remitting; SP, secondary progressive; PP, primary progressive.
